# Assessing Overall Reproducibility for Large-scale High-throughput MRI-based Association Studies

**DOI:** 10.1101/2020.08.18.253740

**Authors:** Zeyu Jiao, Yinglei Lai, Jujiao Kang, Weikang Gong, Liang Ma, Tianye Jia, Chao Xie, Wei Cheng, Andreas Heinz, Sylvane Desrivières, Gunter Schumann, IMAGEN Consortium, Fengzhu Sun, Jianfeng Feng

**Affiliations:** Shanghai Center for Mathematical Sciences, Fudan University, Shanghai, China; Institute of Science and Technology for Brain-inspired Intelligence, Fudan University, Shanghai, China; Key Laboratory of Computational Neuroscience and Brain-Inspired Intelligence (Fudan University), Ministry of Education, China; MOE Frontiers Center for Brain Science, Fudan University, Shanghai, China; Zhangjiang Fudan International Innovation Center; Department of Statistics, The George Washington University, 801 22nd St. NW, Washington, DC 20052, USA; Quantitative and Computational Biology Department, University of Southern California, 1050 Childs Way, Los Angeles, CA, 90089, USA; Department of Computer Science, University of Warwick, Coventry CV4 7AL, UK; School of Life Science and the Collaborative Innovation Center for Brain Science, Fudan University, Shanghai, China; Centre for Functional MRI of the Brain (FMRIB), Nuffield Department of Clinical Neurosciences, Welcome Centre for Integrative Neuroimaging, University of Oxford, Oxford, UK; Key Laboratory of Zoological Systematics and Evolution, Institute of Zoology, Chinese Academy of Sciences, Beijing 100101, China; Centre for Population Neuroscience and Precision Medicine (PONS), Institute of Psychiatry, Psychology & Neuroscience, SGDP Centre, King’s College London, UK; Department of Psychiatry and Psychotherapy CCM, Charité – Universitätsmedizin Berlin, corporate member of Freie Universität Berlin, Humboldt-Universität zu Berlin, and Berlin Institute of Health, Berlin, Germany; PONS Research Group, Department of Psychiatry and Psychotherapy, Campus Charite Mitte, Humboldt University, Berlin, Germany

**Keywords:** Reproducibility, Association studies, MRI (magnetic resonance imaging), Sample size, Heterogeneity

## Abstract

Magnetic Resonance Imaging (MRI) technology has been increasingly used in large-scale association studies. Reproducibility of statistically significant findings generated by MRI-based association studies, especially structural MRI (sMRI) and functional MRI (fMRI), has been recently heavily debated. However, there is still a lack of overall reproducibility assessment for MRI-based association studies. It is also crucial to elucidate the relationship between overall reproducibility and sample size in an experimental design. In this study, we proposed an overall reproducibility index for large-scale high-throughput MRI-based association studies. We performed the overall reproducibility assessments for several recent large sMRI/fMRI databases and observed satisfactory overall reproducibility. Furthermore, we performed the sample size evaluation for the purpose of achieving a desirable overall reproducibility. Additionally, we evaluated the overall reproducibility of GMV changes for UKB vs. PPMI and UKB vs. HCP. We demonstrated that both sample size and some experimental factors play important roles in the overall reproducibility for different experiments. In summary, a systematic assessment of overall reproducibility is fundamental and crucial in the current large-scale high-throughput MRI-based research.

## 1. Introduction

Magnetic Resonance Imaging (MRI) technology has been widely used in neuroscience (Poldrack and Gorgolewski, 2014). It enables us to conduct experiments on grey matter volume (GMV) changes (structure MRI), task-free scanning (resting state fMRI) and task-based studies (task fMRI) (Ashburner and Friston, 2000; Logothetis, 2008; Snyder and Raichle, 2012). Statistical methods for association analyses in these experiments have been frequently performed. Reproducibility, or results reproducibility (Goodman et al., 2016) for MRI-based association studies has recently received a significant attention. Criticisms have been raised to the phenomena that some MRI-based findings are only modestly reproducible and that some results could be interpreted as inflated or spurious (Anonymous, 2017; Bennett and Miller, 2010; Botvinik-Nezer et al., 2020; Eklund et al., 2016). These debates were mostly on the reproducibility of novel discoveries (i.e. findings with statistical significance). To our acknowledge, there is a lack of investigation on the overall reproducibility in MRI-based association studies. A satisfactory overall reproducibility can also provide us with an adequate confidence in MRI-based research outcomes.

Overall reproducibility can be interpreted as the level of concordance among large-scale association analysis results (i.e. z-score). Accordingly, a mixture model based approach has been proposed to conduct hypothesis testing on the overall reproducibility of mass spectrometry studies (Lai et al., 2007). Recently, a Bayesian modeling approach has also been proposed to address the overall irreproducibility of genome-wide association studies or transcriptome-wide association studies (Zhao et al., 2020). For MRI-based association studies, there is still a lack of systematic evaluation of overall reproducibility.

In this study, we proposed an overall reproducibility index for assessing the overall reproducibility of MRI-based association studies. To evaluate the performance of our proposed overall reproducibility index in MRI-based association studies, we first present a comprehensive simulation study which is designed based on UK Biobank structure MRI data (Alfaro-Almagro et al., 2018; Sudlow et al., 2015b). Then, we present overall reproducibility assessments of GMV-related human behavior, brain task state activation and connectivity-behavior studies. Furthermore, with a desirable overall reproducibility requirement, we present the related sample size calculations in various MRI-based study scenarios. These can be achieved with UK Biobank structure and resting-state functional MRI data and IMAGEN task functional MRI data (Bossier et al., 2020; Schumann et al., 2010b). Our sample size calculation results could provide a useful guidance in the related MRI study planning. Moreover, we demonstrate that the overall reproducibility between two independent MRI databases can also be evaluated.

## 2. Materials and Methods

### 2.1 Study participants

#### 2.1.1 UK Biobank

UK Biobank (Alfaro-Almagro et al., 2018; Sudlow et al., 2015b) is a prospective epidemiological resource gathering extensive questionnaires, physical and cognitive measures and biological samples (including genotype), in a cohort of 500000 participants (Sudlow et al., 2015a). Participants which years of age between 40-69 at baseline recruitment consent to access to their full health records from the UK National Health Service, enabling researchers to relate phenotypic measures to long-term health outcomes. They also provided blood, urine and saliva samples, which were stored in such a way as to allow many different types of assay to be performed (for example, genetic, proteomic and metabolomics analyses). In 2014, UK Biobank began the process of inviting back 100000 of the original volunteers for brain, heart and body imaging. The initial release of 10000 UK Biobank imaging and behavioral measures data was used in our manuscript and more details are available online (Alfaro-Almagro et al., 2018).

#### 2.1.2 IMAGEN

One thousand five hundred and six adolescents (mean age = 14.44 y old; SD = 0.42; range = 12.88–16.44 y old) from the baseline assessment of the IMAGEN (Bossier et al., 2020; Schumann et al., 2010b) sample with complete data in fMRI and behavioral measurements were included in the analyses. Detailed descriptions of this study have previously been published (Schumann et al., 2010a).

#### 2.1.3 PPMI

The Parkinson Progression Marker Initiative (PPMI) is a comprehensive observational, international, multi-center study designed to identify PD progression biomarkers both to improve understanding of disease etiology and course and to provide crucial tools to enhance the likelihood of success of PD modifying therapeutic trials. The PPMI cohort will comprise 400 recently diagnosed PD and 200 healthy subjects followed longitudinally for clinical, imaging and biospecimen biomarker assessment using standardized data acquisition protocols at twenty-one clinical sites. We only use the healthy subjects in this cohort and more details see previously publication (Marek et al., 2011).

#### 2.1.4 HCP

The WU-Minn HCP consortium (Van Essen et al., 2012) aims to characterize human brain in a population of 1200 healthy adults and to enable detailed comparisons between brain circuits, behavior, and genetics at the level of individual subjects. Here, we use the T1 data form this project and more details they were initially reported (Van Essen et al., 2013).

### 2.2 MRI acquisition

#### 2.2.1 UK biobank

Details of the image acquisition in UK Biobank are also available online (Alfaro-Almagro et al., 2018). Magnetic resonance imaging (MRI) was performed using a Siemens Skyra 3T running VD13ASP4 (Siemens Healthcare, Erlangen, Germany) with a Siemens 32-channel RF receive head coil. The T1 structural protocol is acquired at 1mm isotropic resolution using a three-dimensional (3D) MPRAGE acquisition, with inversion and repetition times optimized for maximal contrast. The superior-inferior field-of-view is large (256 mm), at little cost, in order to include reasonable amounts of neck/mouth, as those areas will be of interest to some researchers (for example, in the study of sleep apnea).

Resting-state fMRI use the same acquisition parameters, with 2.4-mm spatial resolution and TR = 0.735 s, with multiband acceleration factor. A ‘single band’ reference image (without the multiband excitation, exciting each slice independently) is acquired that has higher tissue-type image contrast; this is used as the target for motion correction and alignment. For both databases, the raw data are corrected for motion and distortion and high-pass filtered to remove temporal drift.

#### 2.2.2 IMAGEN

Structural MRI and fMRI data were acquired at eight IMAGEN assessment sites with 3-T MRI scanners of different manufacturers (Siemens, Philips, General Electric, and Bruker). The scanning variables were specially chosen to be compatible with all scanners. The same scanning protocol was used in all cites. In brief, high-resolution T1-weighted 3D structural images were acquired for anatomical localization and coregistration with the functional time series. BOLD functional images were acquired with a gradient echo, echo planar imaging sequence. 300 vol were acquired for each participant, and each volume consisted of 40 slices aligned to the anterior commission/posterior commission line (2.4-mm slice thickness and 1-mm gap). The echo time was optimized (echo time = 30 ms; repetition time = 2200 ms) to provide reliable imaging of subcortical areas. (More details for different task see **Supplementary**)

#### 2.2.3 PPMI

In this research, we use the MRI data acquired by the PPMI study, in which a T1-weighted, 3D sequence (i.e., MPRAGE) is acquired for each subject using 3T SIEMENS MAGNETOM TrioTim syngo scanners. This gives us 374 PD and 169 NC scans. The T1-weighted images were acquired for 176 sagittal slices, with the following parameters: repetition time (TR) = 2300ms, echo time (TE) = 2.98ms, flip angle = 9°, and voxel size = 1 × 1 × 1mm^3^.

#### 2.2.4 HCP

The Human Connectome Project (HCP) provides a unique, open source, large-scale collection of about 1200 human head T1 image datasets and we employed 413 healthy subjects which have no clear family related (age-range: 22-36 years) in our study. All HCP imaging data were acquired on a Siemens Skyra 3T scanner with a customized SC72 gradient insert. T1w 3D MPRAGE were acquired with TR = 2400 ms, TE = 2.14 ms, TI = 1000 ms, flip angle =8deg, FOV =224×224, 0.7 mm isotropic voxel, bandwidth = 210 Hz/px, iPAT = 2, Acquisition time = 7:40 (min:sec).

### 2.3 MRI Preprocessing

The rs-fMRI data are preprocessed using standard volume-based fMRI pipeline. For each subject, the preprocessing steps include: motion correction (FSL mcflirt), despiking motion artifacts using Brain Wavelet Toolbox (Patel et al., 2014), registering to 3×3×3 mm^2^ standard space by first aligning the functional image to the individual T1 structure image using boundary based registration (Greve and Fischl, 2009) and then to standard space using FSL’s linear and non-linear registration tool (FSL flirt and fnirt), regressing out nuisance covariates including Friston-24 parameters, white matter signal, cerebrospinal fluid signal, band-pass filtering (0.01-0.1 Hz) using AFNI (3dTproject) and spatial smoothing by a 3D Gaussian kernel (FWHM= 6 mm). All the images are manually checked to ensure successful preprocessing and insure the mean FD Power not greater than 0.5. After above preprocessing, a large sample size imaging and behavioral measures data which contain 8273 subjects have been used in this study.

T1 data were preprocessed with the voxel-based morphometry (VBM) by using the VBM8 toolbox based on the Statistical Parametric Mapping package (SPM). Firstly, all structural MRI data were manually corrected and divided into grey matters, white matters and cerebrospinal fluid. Secondly, the grey matter images were aligned to a nonlinear deformation field and normalized to Montreal Neurological Institute (MNI) space by using the templates which were created by DARTEL tool. Finally, the normalized images were all smoothed with a full-width at half-maximum (FWHM) 6-mm Gaussian kernel for further analysis. After above procedures, the grey matter images (voxel size: 3×3×3 mm^2^) were obtained for 9850 subjects.

Task-fMRI data were analyzed with SPM. Spatial preprocessing included slice time correction to adjust for time differences caused by multi-slice imaging acquisition, realignment to the first volume in line, nonlinearly warping to the Montreal Neurological Institute space [based on a custom echo planar imaging template (53×63×46 mm^3^ voxels) created out of an average of the mean images of 400 adolescents], resampling at a resolution of 3×3×3 mm^3^, and smoothing with an isotropic Gaussian kernel of 5-mm FWHM.

At the first level of analysis, changes in the BOLD response for each subject were assessed by linear combinations at the individual subject level, for each experimental condition (e.g. reward anticipation high gain of Monetary Incentive Delay (MID) task), each trial was convolved with the hemodynamic response function to form regressors that account for potential noise variance, e.g. head movement, associated with the processing of reward anticipation. Estimated movement parameters were added to the design matrix in the form of 18 additional columns (three translations, three rotations, three quadratic and three cubic translations, and every three translations with a shift of ±1 TR).

For the MID anticipation phase we contrasted brain activation during ‘anticipation of high win [here signaled by a circle] vs anticipation of no-win [here signaled by a triangle]’; For the emotional faces task (EFT) we contrasted brain activation during ‘viewing Angry Face vs viewing Control [circles]’; For the stop signal task (SST) we contrasted brain activation during ‘successful stop vs successful go’.

### 2.4 Statistical Analyses

#### 2.4.1 Functional Connectivity Association Study

Based on the automated anatomical labeling (AAL2) atlas, there are 120 brain regions. Each resting-state functional magnetic resonance image (rs-fMRI) included 54,885 voxels (Rolls et al., 2015). For each pair of brain regions, the time series were extracted, and the Pearson correlation was calculated for each subject to provide the measure of functional connectivity (FC), followed by Fisher’s z-transformation. The general linear model was used to test the association between the region-wise FC links and a human phenotype or behavior. The effects of age, sex and head motion (mean frame-wise displacement) were regressed out.

#### 2.4.2 Voxel-wise Association Study

We used the general linear model to define the association between a specific human phenotype or behavior and each intracerebral voxel’s gray matter volume, which was included in the automated anatomical labeling (AAL2) atlas (total 54,885 voxels). The effects of age, sex and total intracerebral volume (TIV) were regressed out.

#### 2.4.3 Task fMRI Activation

At the first level of analysis, changes in the BOLD response for each subject were assessed by linear combinations at the individual subject level for each experimental condition, and each trial was convolved with the hemodynamic response function to form regressors that account for potential noise variance (e.g., head movement) associated with the processing of a specific task. Estimated movement parameters were added to the design matrix in the form of 18 additional columns (three translations, three rotations, three quadratic and three cubic translations, and three translations each with a shift of ±1 repetition time). To identify brain activation specific to the task, we contrasted the brain activation patterns between the task status and the control status.

#### 2.4.4 Normal Distribution Quantile-based Transformation

*z*-scores from a normal distribution quantile transformation were used for the analysis (Lai et al., 2007). First, based on an appropriate association analysis (functional connectivity association study, voxel-wise association study or task fMRI activation), we acquired a list of one-sided *P*-values. For each *P*-value *P*, the corresponding *z*-score *z* can be calculated as follows:

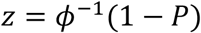

where *ϕ*^−1^(·) is the inverse function of the standard normal cumulative distribution function.

#### 2.4.5 Definition of Overall Reproducibility Index

We firstly consider a nine-component normal-mixture model for the joint distribution of paired *z*-scores [*z*^(1)^, *z*^(2)^] (see above for *z*-score calculation).

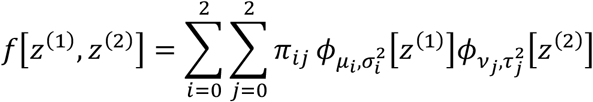

where 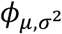 is the normal probability distribution function with mean *μ* and variance *σ*^2^. We use the first component (index 0) to represent the null (no change/correlation) feature component. Then, *μ*_0_ = *v*_0_ = 0 and 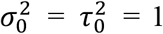. The second and third components (indices 1 and 2) are used to represent negative changes/correlations and positive changes/correlations. Their corresponding parameters (means and variances) will be estimated from the paired *z*-scores with the following constraints: *μ*_1_, *v*_1_ ≤ 0 and *μ*_2_, *v*_2_ ≥ 0. *π*_*ij*_ is the proportion for component *i* in the first association study and component *j* in the second association study, and Σ_*ij*_ *π*_*ij*_ = 1.

This model was termed partial concordance/discordance (PCD) model (Lai et al., 2007; Lai et al., 2009; Lai et al., 2014, 2017). Then, we define an overall reproducibility index based on model parameters, mixture model reproducibility index (*M*^*2*^*RI*) (The illustration of *M*^*2*^*RI* see **Figure 1**):

**Figure 1:**
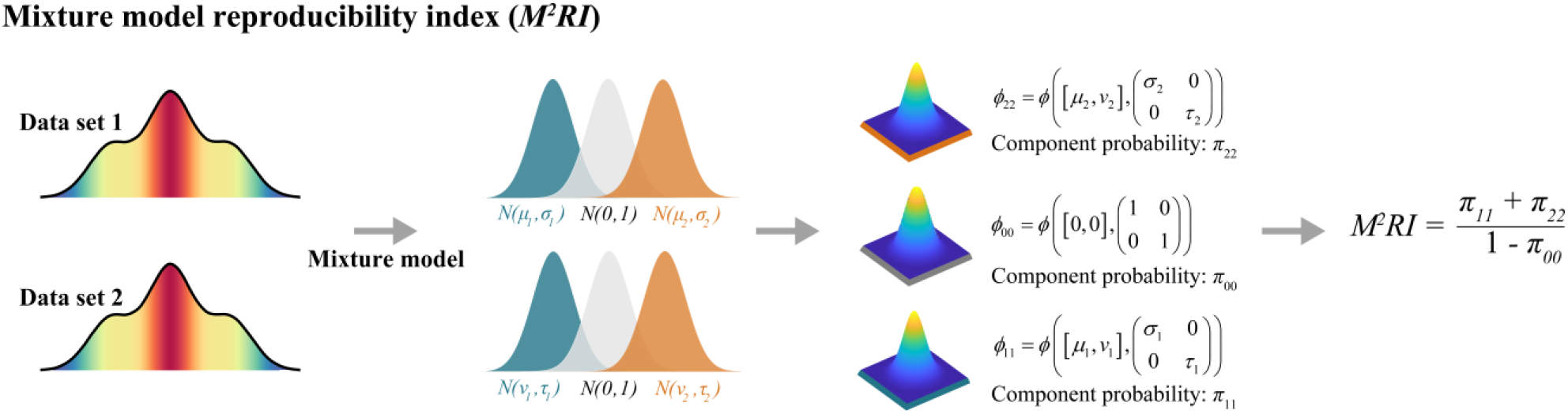
An illustration of *M*^*2*^*RI*. *ϕ*_*i,j*_ is the normal probability distribution function and *π*_*ij*_ is the proportion of features consistent with component *i* in the first association analysis and component *j* in the second association analysis.

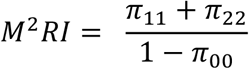

In a recent study (Zhao et al., 2020), two Bayesian models: curved exponential family normal prior model (CEFN) and meta-analysis prior model (META), have been proposed for a similar purpose. In these two models, *π*_*null*_ and *π*_*R*_ were the proportions of null and reproducible signals, respectively. Accordingly, we may define the related overall reproducibility indices:

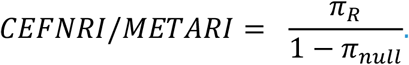

#### 2.4.6 Confidence Intervals of *M*^*2*^*RI*

The confidence intervals (CIs) of *M*^*2*^*RI* can be obtained by bootstrapping paired z-scores (Efron and Tibshirani, 1997; Mclachlan, 1987). For our newly developed overall reproducibility index *M*^*2*^*RI*, a theoretical confidence interval will also be highly useful in practice. Therefore, we have derived the asymptotic theoretical CIs for *M*^*2*^*RI* based on our proposed mixture model (see **Supplementary** for details).

## 3. Results

### 3.1 Overall Reproducibility Index Recovers the True Overall Reproducibility Accurately in the Simulation Study

We conducted a comprehensive simulation study to show the performance of our newly proposed overall reproducibility index. Our simulations were designed based on the gray matter volume (GMV) data in the UK Biobank. Two-sample comparison is a general association analysis scenario in practice, and the overall reproducibility of a large-scale two-sample study is important. Therefore, we partitioned the data randomly into four subsets (referred to as Data 1A, Data 1B, Data 2A and Data 2B). Before the analysis, as a widely considered practical approach, we checked that sex, age, total intracerebral volume (TIV) and total GMV were statistically similar between Data 1A vs. 2A as well as Data 1B vs. 2B (two-sample *t*-test, *P*>0.05). Otherwise, we repeated the random data partition until one passed this similarity requirement. For each feature, there was statistically no differences in distribution between Data 1A vs. 2A nor Data 1B vs. 2B. Then, to generate upward or downward changes, a specified proportion of voxels in a cluster were randomly chosen and 0.0285-0.0855 standard deviations of brain-wise GMV (corresponding to approximately 1-3 effect sizes in *z*-scores) were randomly added to (or subtracted from) the chosen voxels of each subject in Data 1A and Data 1B. This procedure was repeated 1,000 times. For each repetition, we obtained two lists of *z*-scores: one by voxel-wisely comparing Data 1A vs. Data 2A and the other Data 1B vs. Data 2B. *z*-scores were calculated based on the traditional two-sample *t*-test. A pair of *z*-scores were obtained for each voxel. The overall reproducibility between two lists of *z*-scores was assessed by our proposed overall reproducibility index. The following three simulation scenarios were considered.

#### (a) Complete overall reproducibility with a moderate proportion of changes

According to our random data partition, there were statistically no differences between Data 1A vs. 2A nor Data 1B vs. 2B. We modified the 100% of null (no change) to 80% null, 10% upward changes and 10% downward changes as follows. We randomly selected two clusters of voxels, each with 10% of the total voxels. To simulate 10% upward changes, for each voxel in the first cluster of voxels, we randomly added to each subject’s GMV a value equivalent to 1-3 effect sizes in *z*-scores in Data 1A and repeated this in Data 1B so that there were 10% reproducible upward changes. For each voxel in the second cluster of voxels, we randomly subtracted from each subject’s GMV a value equivalent to 1-3 effect sizes in *z*-score in Data 1A and repeated this in Data 1B so that there were 10% reproducible downward changes.

#### (b) Partial reproducibility

We randomly selected four clusters of voxels. There were 15% of the total voxels in each of the first two clusters, and the upward changes and downward changes were simulated according to the description in (a). There were 5% of the total voxels in each of the next two clusters. For each voxel in the third cluster, we randomly added to each subject’s GMV a value equivalent to 1-3 effect sizes in *z*-scores in Data 1A (but not in Data 1B). Then, we had 5% discordant changes (up vs. null). For each voxel in the fourth cluster, we similarly subtracted from each subject’s GMV in Data 1A (but not in Data 1B) so that we had 5% discordant changes (down vs. null).

#### (c) Complete overall reproducibility with a high proportion of changes

Considering that the number of consistent significant results from different studies can vary, we randomly selected two clusters of voxels, each with 20% of the total voxels. The reproducible upward changes (the first cluster) and downward changes (the second cluster) were simulated similarly according to the description in (a).

The results are summarized in **Table 1**. Based on the scenario (a) as complete overall reproducibility with a moderate proportion of changes, the median *M*^*2*^*RI*, *CEFNRI* or *METARI* was 0.915, 0.995 or 0.988, respectively. Furthermore, the related lower- and upper-quartiles (Q1-Q3) was 0.738-0.995, 0.993-0.996 or 0.951-0.993, respectively. It was reasonable to conclude that the assessed overall reproducibility could be up to the true overall reproducibility which is 100%. Based on the scenario (b) as a partial reproducibility (75%), the median *M*^*2*^*RI*, *CEFNRI* or *METARI* was 0.769, 0.988 or 0.826 when the related lower- and upper-quartiles (Q1-Q3) was 0.685-0.849, 0.955-0.995 or 0.691-0.909, respectively. Based on the scenario (c) as complete overall reproducibility with a high proportion of changes, the median *M*^*2*^*RI*, *CEFNRI* or *METARI* reached 0.960, 0.998 or 0.994 with the lower- and upper-quartiles (Q1-Q3) 0.873-0.999, 0.997-0.998 or 0.973-0.997, respectively. It was also reasonable to conclude that the assessed overall reproducibility could be up to the true overall reproducibility which is 100%.

**Table 1:** The performance of overall reproducibility index in three simulation analysis scenarios. For each simulation analysis scenario, the true reproducibility index is shown in the table. The simulation and evaluation were repeated 1000 times to obtain the median, the lower and upper-quartiles (Q1-Q3) for assessed overall reproducibility. (For more details, please see section *Overall Reproducibility Index Recovers the True Overall Reproducibility Accurately in the Simulation Study*.)

| ***Overall Reproducibility Index*** | ***Assessed Overall Reproducibility*** |
| --- | --- |
| *Simulation (a)* | *Median (Q1-Q3)* |
| ***100% reproducibility*** |  |
| *M<sup>2</sup>RI* | 0.9150 (0.7378 – 0.9950) |
| *CEFNRI* | 0.9951 (0.9932 – 0.9961) |
| *METARI* | 0.9879 (0.9506 – 0.9933) |
| *Simulation (b)* | *Median (Q1-Q3)* |
| ***75% reproducibility*** |  |
| *M<sup>2</sup>RI* | 0.7688 (0.6850 – 0.8488) |
| *CEFNRI* | 0.9875 (0.9549 – 0.9948) |
| *METARI* | 0.8255 (0.6911 – 0.9092) |
| *Simulation (c)* | *Median (Q1-Q3)* |
| ***100% reproducibility*** |  |
| *M<sup>2</sup>RI* | 0.9597 (0.8731 – 0.9986) |
| *CEFNRI* | 0.9980 (0.9973 – 0.9983) |
| *METARI* | 0.9939 (0.9734 – 0.9966) |

### 3.2 Overall Reproducibility of Large-scale MRI-based Association Studies

To investigate the overall reproducibility of large-scale MRI-based association analysis in the data collected for studying human phenotypes/behaviors and task state activations, as well as the brain structure and function, we split each study cohort into two subsets (referred to as Group 1 and Group 2 based on the order of subject number) with (approximately) the same sample sizes. For the resting-state functional connectivity (RSFC) data, the sample sizes of the two subsets were 4,136 and 4,137 for analyzing sex as phenotype vs. RSFC; the sample sizes of the two subsets were 4,131 and 4,131 for analyzing body mass index (BMI) as phenotype vs. RSFC (as there were missing BMI observations). A general linear model was constructed with sex phenotype as the response in each subset, with age and mean FD adjusted as covariates (hereafter referred to as Sex as phenotype vs. RSFC and BMI as phenotype vs. RSFC; see **Figure 2c** and **Figure 2d** for the paired *z*-scores). For the GMV data, the sample sizes of the two subsets were 4,925 and 4,925, respectively. A general linear model was also constructed with sex phenotype as the response in each subset, with age and TIV adjusted as covariates (hereafter referred to as Sex as phenotype vs. GMV; see **Figure 2a** for the paired *z*-scores). For the task-related activation data, the sample sizes of the two subsets were 772 and 772, respectively. Student’s *t*-test was used to evaluate the activation of the monetary incentive delay (MID) task, one of the most common tasks in fMRI studies (this activity is hereafter referred to as Activation in the MID task; see **Figure 2b** for the paired *z*-scores). For each paired *z*-scores, an overall diagonal pattern can be clearly observed. Different paired *z*-scores variation patterns can also be observed for different analysis scenarios, which implies different mixtures of no-change related (null) *z*-scores and upward/downward-change related (non-null) *z*-scores.

**Figure 2:**
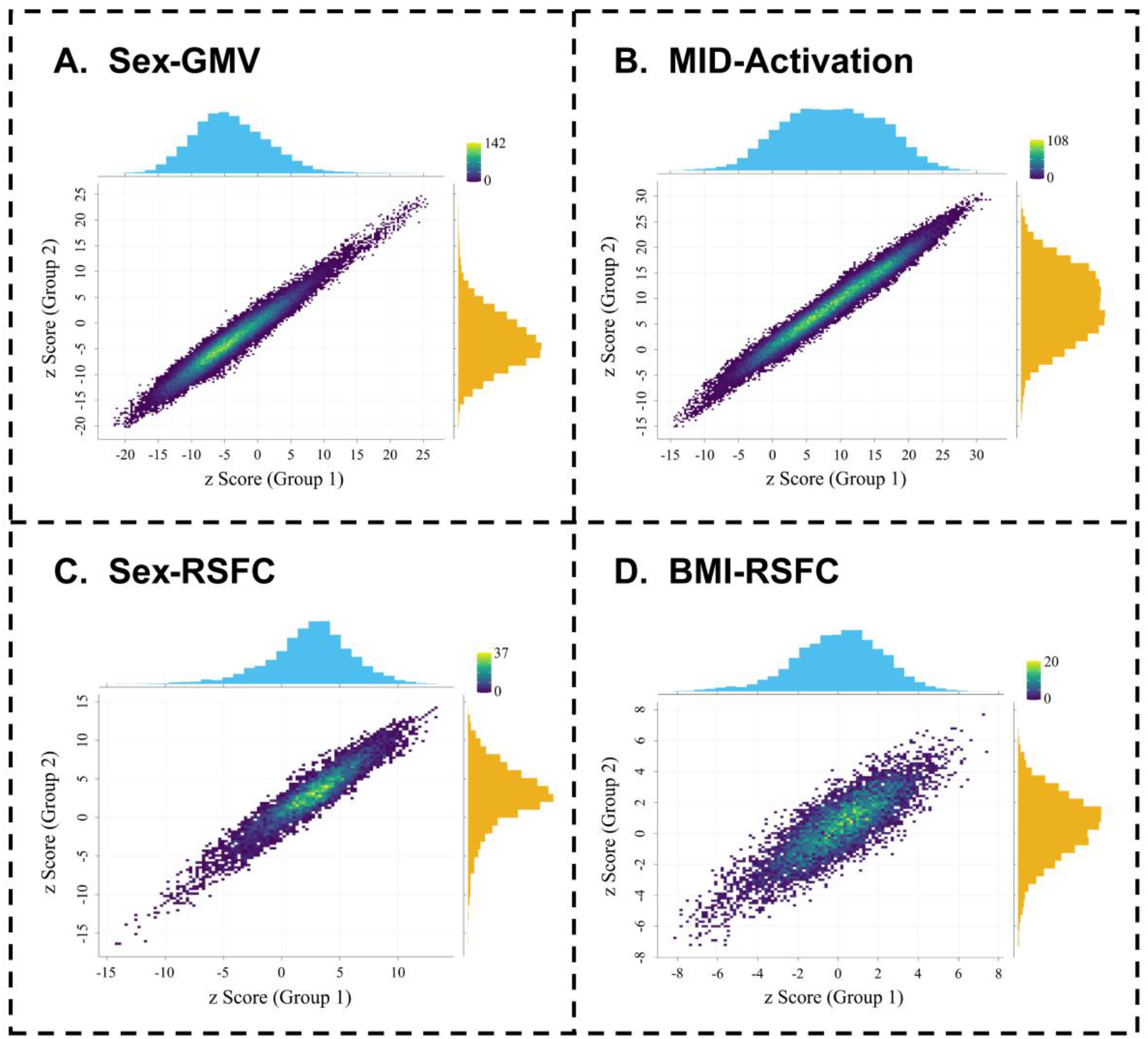
Paired *z*-scores from four association analysis scenarios. (A) Sex as phenotype vs. GMV in UK Biobank data. (B) MID task activation in IMAGEN data. (C) Sex as phenotype vs. RSFC in UK Biobank data. (D) BMI as phenotype vs. RSFC in UK Biobank.

Overall reproducibility index was used to evaluate the overall reproducibility based on the paired *z*-scores in **Figure 2**. The results are shown in **Table 2**. We bootstrapped the paired *z*-score to construct the related 95% confidence intervals (CIs) and we also calculated the asymptotic theoretical 95% CIs for *M*^*2*^*RI*. For Sex as phenotype vs. RSFC, the *CEFNRI* or *METARI* was 0.9993 or 0.9992, respectively. Furthermore, *M*^*2*^*RI* was close to one, which also suggested an ideal overall reproducibility. Its asymptotic theoretical 95% CI was above 0.98. For BMI as phenotype vs. RSFC, *CEFNRI* or *METARI* was 0.9986 or 0.9984, respectively. *M*^*2*^*RI* was still close to one, and its asymptotic theoretical 95% CIs were above 0.97. For Sex as phenotype vs. GMV, *CEFNRI* or *METARI* was 0.9996 or 0.9995, respectively. *M*^*2*^*RI* was again nearly one and both 95% CIs were nearly ideal. For activation in the MID task, *CEFNRI* or *METARI* was 0.9997 or 0.9997, respectively. *M*^*2*^*RI* was still nearly one and both 95% CIs were again nearly ideal.

**Table 2:** Overall reproducibility assessment of four association analysis scenarios. For each MRI-based association analysis scenario, the assessed overall reproducibility is shown in the table. We obtained 95% confidence intervals (CIs) based on bootstrapping the paired *z*-scores for *M*^*2*^*RI*. The asymptotic theoretical 95% CIs for *M*^*2*^*RI* was also presented. (For more details, please see section *Overall Reproducibility of Large-scale MRI-based Association Studies*.)

| **Overall Reproducibility Index** | **Assessed Overall Reproducibility** |
| --- | --- |
| *Sex as phenotype vs. RSFC* |  |
| $M^2RI$ (95% CIs) | 0.9999996 (0.9999993 – 0.9999998) |
| $M^2RI$ (asymptotic theoretical 95% CIs) | 0.9999996 (0.9807 – 1) |
| CEFNRI | 0.9993 |
| METARI | 0.9992 |
| *BMI as phenotype vs. RSFC* |  |
| $M^2RI$ (95% CIs) | 0.9999934 (0.9999904 – 0.9999955) |
| $M^2RI$ (asymptotic theoretical 95% CIs) | 0.9999934 (0.9782 – 1) |
| CEFNRI | 0.9986 |
| METARI | 0.9984 |
| *Sex as phenotype vs. GMV* |  |
| $M^2RI$ (95% CIs) | 0.99999995 (0.99999993 – 0.99999997) |
| $M^2RI$ (asymptotic theoretical 95% CIs) | 0.99999995 (0.9955 – 1) |
| CEFNRI | 0.9996 |
| METARI | 0.9995 |
| *Activation in MID task* |  |
| $M^2RI$ (95% CIs) | 0.99999991 (0.99999989 – 0.99999994) |
| $M^2RI$ (asymptotic theoretical 95% CIs) | 0.99999991 (0.9968 – 1) |
| CEFNRI | 0.9997 |
| METARI | 0.9997 |

### 3.3 Sample Size Needed to Achieve a Desirable Overall Reproducibility

Sample size calculation is crucial in experimental designs. When designing a large-scale association analysis, one may ask what sample size is required to achieve a desirable overall reproducibility requirement. For a comprehensive understanding of sample size requirements in different large-scale association analysis scenarios, we conducted a large resampling-based simulation study. For a study cohort presented in **Table 2**, we selected a phenotype available in the study as response. Then, we randomly selected subjects from the cohort to construct two subsets with a given sample size for each subset. This procedure is appropriate for *M*^*2*^*RI*. For each given sample size, we repeated the resampling and *M*^*2*^*RI* calculation 1,000 times.

We evaluated Pr(*M*^*2*^*RI* > 0.8) empirically for each given sample size. Then, we could obtain the minimum sample size to achieve Pr(*M*^*2*^*RI* > 0.8) > 0.8 in each analysis scenario. (In addition to 0.8, other values could be certainly considered, and it is not necessary to always set both values to 0.8.) The results for different analysis scenarios are summarized in **Figure 3** and **Table 3**. We assessed the minimum sample size for *M*^*2*^*RI*. For different response phenotypes in the task-related functional MRI data, the minimum sample size was only approximately 20 to 30. For the GMV data, a sample size of approximately 120 was required when the response was the sex phenotype; a sample size of 70 was required when the response was the age phenotype and a sample size of 300 was required when the response was the BMI phenotype. However, for different response phenotypes in the RSFC data, the results were clearly different. Approximately 200 or 300 were required when the response was the age or sex phenotype, respectively. When the response was BMI, the minimum sample size increased to a very large value (More than 800. When the estimated minimum sample size *n* close to the sample size of the whole dataset *N* (e.g. *n* is more than 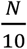), the minimum sample size estimation could be inaccurate as the sample duplication problem in resampling-based simulation study).

**Table 3:** *M*^*2*^*RI* based sample size calculations in different MRI association analysis scenarios. The minimum sample size to achieve Pr(*M*^*2*^*RI* > 0.8) > 0.8 is presented for each large-scale association analysis scenario. For the RSFC data, sex, age or BMI was considered as phenotype. For the GMV data, sex, age or BMI was considered as phenotype. For the fMRI data in task activation, MID, SST or EFT task was considered. (For more details, see section *Sample Size Needed to Achieve a Desirable Overall Reproducibility*.)

| *MRI Study* | *Minimum*<br>*Sample Size* | *MRI Study* | *Minimum*<br>*Sample Size* | *MRI Study* | *Minimum*<br>*Sample Size* |
| --- | --- | --- | --- | --- | --- |
| *RSFC* |  | *GMV* |  | *Task fMRI* |  |
| *Sex* | **300** | *Sex* | **120** | *MID* | **30** |
| *Age* | **200** | *Age* | **70** | *SST* | **20** |
| *BMI* | **More than**<br>**800** | *BMI* | **300** | *EFT* | **30** |

**Figure 3:**
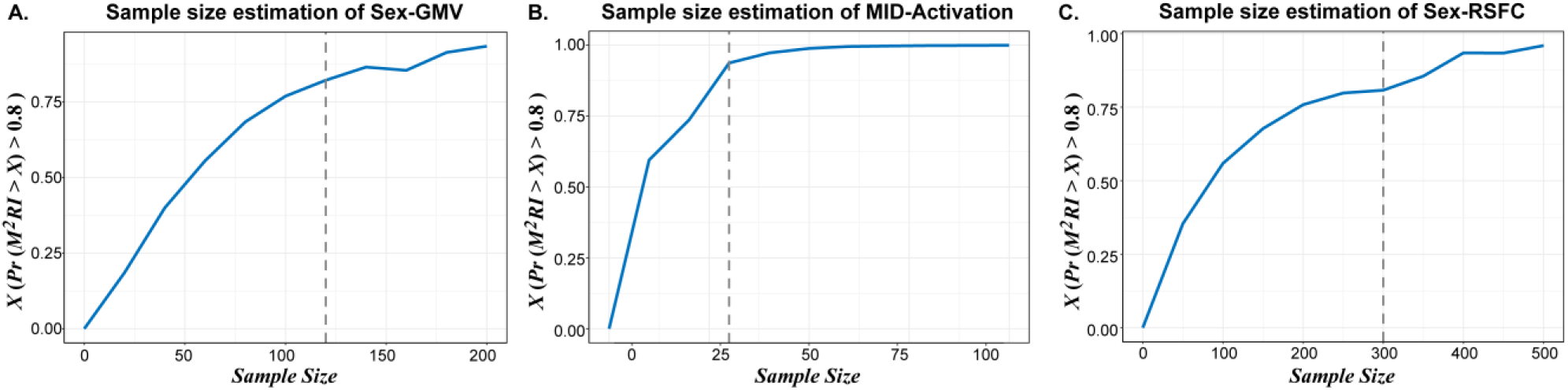
Sample size calculations for three association analysis scenarios in the UK Biobank or IMAGEN data. In each plot, “*X* (*Pr*(*M*^*2*^*RI* > *X*) > 0.8)” means the minimal *M*^*2*^*RI* in top 80% resampled repetitions (800 resampled repetitions in total 1,000 resampled repetitions). The vertical dashed line indicates the minimum sample size for Pr (*M*^*2*^*RI* > 0.8) > 0.8. (A) Sex as phenotype vs. GMV in the UK Biobank data. (B) MID task activation in the IMAGEN data. (C) Sex as phenotype vs. RSFC in the UK Biobank data.

### 3.4 Application: Overall Reproducibility Evaluation of GMV Change for UKB vs. PPMI and UKB vs. HCP

As an application of overall reproducibility index, we considered two MRI databases: PPMI and UK Biobank cohorts. For the PPMI database, there were 136 normal subjects of age from 45 to 79. As the UK Biobank cohort is much larger, we performed a sample matching based on age and sex for this analysis. Seven age groups of 45-49, 50-54, etc. (5-year intervals) were considered. For each age group, from the UK Biobank cohort, we randomly selected the same number of female/male subjects as that in the PPMI cohort. A total of 136 subjects were randomly selected from the UK Biobank cohort. Then, for both databases, we calculated the *z*-scores for the age phenotype as response vs. GMV based on a general linear model with the adjustments for sex and TIV. This was repeated 1,000 times and we obtained 1,000 lists of paired *z*-scores.

As another application of overall reproducibility index, we considered the HCP and UK Biobank cohorts. For the HCP data set, there were 413 subjects of age from 22 to 36. Then, it was not feasible to match the corresponding age ranges in the UK Biobank data because this age range was not available in the UK Biobank. We still performed a sample matching based on sex. From the UK Biobank cohort, we randomly selected the same number of female/male subjects as that in the HCP cohort. A total of 413 subjects were randomly selected from the UK Biobank cohort. Then, for both databases, we calculated the *z*-scores for the sex phenotype as response vs. GMV based on a general linear model with the adjustments of age and TIV. This was repeated 1,000 times and we obtained 1,000 lists of paired *z*-scores.

For each list of paired z-scores, we applied overall reproducibility index to assess the related overall reproducibility (see **Table 4** for results). For the PPMI and UK Biobank databases, the median *M*^*2*^*RI*, *CEFNRI* or *METARI* was 0.99993,0.9974 or 0.8642 with the lower- and upper-quartiles (Q1-Q3) 0.99964-0.99998, 0.9967-0.9979 or 0.7404-0.9625, respectively. It was reasonable to conclude that both databases were ideally overall reproducible in term of large-scale association analysis with age as phenotype. For the HCP and UK Biobank databases, the median *M*^*2*^*RI*, *CEFNRI* or *METARI* was only 0.6378,0.4377 or 0.0420, with the lower- and upper-quartile (Q1-Q3) 0.5747-0.7032, 0.3922-0.4838 or 0.0263-0.0656, respectively. As the age ranges for both databases were clearly different, it was also reasonable to observe a relatively low overall reproducibility for this analysis.

**Table 4:** Overall reproducibility index for comparing two databases. Overall reproducibility index applications in comparing the large-scale association analysis results from two closely related databases. Median with the range of interquartile (Q1-Q3) are shown in the table. For the MRI databases PPMI vs. UK Biobank, the analysis was based on age as phenotype vs. GMV. For the MRI databases HCP vs. UK Biobank, the analysis was based on sex as phenotype vs. GMV. (For more details, please see section *Application: Overall Reproducibility Evaluation of GMV Change for UKB vs. PPMI and UKB vs. HCP*.)

| **Application** | **Assessed overall reproducibility** |
| --- | --- |
| PPMI vs. UKB (Sample size = 136) | *Median (Q1-Q3)* |
| $M^2RI$ | 0.99993 (0.99964 – 0.99998) |
| *CEFNRI* | 0.9974 (0.9967 – 0.9979) |
| *METARI* | 0.8642 (0.7404 – 0.9625) |
| HCP vs. UKB (Sample size = 413) | *Median (Q1-Q3)* |
| $M^2RI$ | 0.6378 (0.5747 – 0.7032) |
| *CEFNRI* | 0.4377 (0.3922 – 0.4838) |
| *METARI* | 0.0420 (0.0263 – 0.0656) |

## 4. Discussion

Reproducibility of discoveries based on large-scale data has received a significant attention in recent years. However, there are still a lack of overall reproducibility evaluation in neuroimaging studies (Anonymous, 2017; Botvinik-Nezer et al., 2020; Eklund et al., 2016; Poldrack, 2019). To address this need, we proposed a mixture model based overall reproducibility index, and we discussed its relationship to a recently proposed irreproducibility quantity (Zhao et al., 2020). Through a comprehensive simulation study, we demonstrated the advantages of overall reproducibility index for an accurate overall reproducibility assessment in large-scale MRI-based association studies. We also demonstrated satisfactory overall reproducibility achieved from UK Biobank or IMAGEN based large-scale association analyses. Then, we evaluated the sample size necessary for achieving a desirable overall reproducibility, which is essential in a design of experiment. Moreover, we evaluated the overall reproducibility of GMV changes for UKB vs. PPMI and UKB vs. HCP. We still observed satisfactory results based on UKB vs. PPMI because of their highly similar experimental factors. However, we could not observe satisfactory results based on UKB vs. HCP because of their experimental factors could not be entirely similar. Therefore, the impact from some experimental factors plays an important role when the overall reproducibility is evaluated.

Our newly proposed overall reproducibility index was derived from the combination of parameters in a mixture model. This index is close related to the irreproducibility quantity *ρ*_*IR*_ that has been proposed in a previous study (Zhao et al., 2020). Although the overall reproducibility index can be interpreted as 1 − *ρ*_*IR*_, their related statistical models are different.

We conducted a comprehensive sample size calculation for several recent large sMRI/fMRI databases. According to our results, an adequate sample size is necessary to report a reliable overall reproducibility assessment. Additionally, the sample size requirement is closely related to the strength of associations, which depends largely on the signal-to-noise ratio of response outcome (e.g., phenotypes) and predictors (e.g., MRI signal). Therefore, the impact of different phenotypes, predictor data types, and technology platforms should all be considered in the study of overall reproducibility assessment. These results are well illustrated in our results. To achieve the desirable overall reproducibility, the required sample size for a task fMRI study is clearly lower than that for a GMV study, which is clearly lower than that for an RSFC study. For a GMV study, the required sample size for the age phenotype as response is clearly lower than that for the sex phenotype as response. For an RSFC study, the required sample size for the BMI phenotype as response is much larger. These results are consistent with our expectations. The data signal-to-noise ratios from a task fMRI study are usually clearly large, and the data signal-to-noise ratios from the GMV study are usually comparably larger than those from the RSFC study. The phenotype signal-to-noise ratio of BMI is clearly smaller than that of sex or age phenotype. Therefore, our results are highly illustrative and informative for planning the sample size for a large-sale high-throughput MRI-based association study.

As data pooling or meta-analysis is frequently considered in practice, the evaluation of overall reproducibility between two closely related studies is crucial. Such an analysis allows us to understand the heterogeneity among different studies. It also allows us to address the generalizability of a large-scale association analysis. The relationship between the overall reproducibility and the heterogeneity among different experiments was also discussed (Zhao et al., 2020).

We have demonstrated that our proposed index is useful for the overall reproducibility assessment of large databases. It is still necessary to further develop novel and useful tools for data with relatively small sample sizes. Statistically, when the sample size is relatively small, it is difficult to fit the *z*-scores with a simple model. As our future research endeavor, we will investigate other approaches so that the overall reproducibility assessment can be achieved for data with a relatively small sample size. We believe that these efforts will also help improve the current approach for data with a relatively large sample size. Moreover, overall reproducibility index can also be applied to brain-wide association study (BWAS) (Cheng et al., 2015a; Cheng et al., 2015b; Gong et al., 2018) and other types of large-scale high-throughput association study. We would point out that the overall reproducibility assessment of BWAS can be computationally time consuming. As the number of features (voxel-wise FCs) is significantly large, we will direct our future research endeavors toward how to conduct such an analysis more effectively and efficiently.

## 5. Conclusions

The reproducibility of large-scale high-throughput MRI-based research discoveries has been recently debated. There is still a lack of effective and efficient overall reproducibility assessment in neuroimaging experiments. Therefore, we have developed an overall reproducibility index for assessing the overall reproducibility in large-scale MRI-based association studies. For several recent well-known large-sample studies, we have evaluated the sample size to achieve a desirable overall reproducibility requirement. Our study provides a scientific contribution to the measurement of overall reproducibility that is fundamental and crucial in the current large-scale high-throughput MRI-based research.

## Acknowledgements

This work received support from the following sources: National Key R&D Program of China (No. 2019YFA0709502), National Key R&D Program of China (No.2018YFC1312904), the 111 Project (NO.B18015), the key project of Shanghai Science &Technology (No.16JC1420402), Shanghai Municipal Science and Technology Major Project (No.2018SHZDZX01), ZJ Lab, and Shanghai Center for Brain Science and Brain-Inspired Technology, National Key R&D Program of China (No 2018YFC1312900), National Key R&D Program of China (No. 2019YFA0709501), National Natural Science Foundation of China (NSFC 91630314, 81801773), National Natural Science Foundation of China (No. 11971459, 81673833), the European Union-funded FP6 Integrated Project IMAGEN (Reinforcement-related behaviour in normal brain function and psychopathology) (LSHM-CT-2007-037286), the Horizon 2020 funded ERC Advanced Grant ‘STRATIFY’ (Brain network based stratification of reinforcement-related disorders) (695313), ERANID (Understanding the Interplay between Cultural, Biological and Subjective Factors in Drug Use Pathways) (PR-ST-0416-10004), BRIDGET (JPND: BRain Imaging, cognition Dementia and next generation GEnomics) (MR/N027558/1), Human Brain Project (HBP SGA 2, 785907), the FP7 project MATRICS (603016), the Medical Research Council Grant ‘c-VEDA’ (Consortium on Vulnerability to Externalizing Disorders and Addictions) (MR/N000390/1), the National Institute for Health Research (NIHR) Biomedical Research Centre at South London and Maudsley NHS Foundation Trust and King’s College London, the Bundesministerium für Bildung und Forschung (BMBF grants 01GS08152; 01EV0711; Forschungsnetz AERIAL 01EE1406A, 01EE1406B), the Deutsche Forschungsgemeinschaft (DFG grants SM 80/7-2, SFB 940, TRR 265, NE 1383/14-1), the Medical Research Foundation and Medical Research Council (grants MR/R00465X/1 and MR/S020306/1), the National Institutes of Health (NIH) funded ENIGMA (grants 5U54EB020403-05 and 1R56AG058854-01), the Human Brain Project (HBP SGA 2). Further support was provided by grants from ANR (project AF12-NEUR0008-01 - WM2NA, and ANR-12-SAMA-0004), the Fondation de France, the Fondation pour la Recherche Médicale, the Mission Interministérielle de Lutte-contreles-Drogues-et-les-Conduites-Addictives (MILDECA), the Assistance-Publique-Hôpitaux-de-Paris and INSERM (interface grant), Paris Sud University IDEX 2012; the National Institutes of Health, Science Foundation Ireland (16/ERCD/3797), U.S.A. (Axon, Testosterone and Mental Health during Adolescence; RO1 MH085772-01A1) and by NIH Consortium grant U54 EB020403, supported by a cross-NIH alliance that funds Big Data to Knowledge Centres of Excellence. The funders had no role in the study design, data collection and analysis, decision to publish or preparation of the manuscript.

## References

Anonymous, 2017. Fostering reproducible fMRI research. Nature Neuroscience 20, 298–298.

Alfaro-Almagro, F., Jenkinson, M., Bangerter, N.K., Andersson, J.L.R., Griffanti, L., Douaud, G., Sotiropoulos, S.N., Jbabdi, S., Hernandez-Fernandez, M., Vallee, E., Vidaurre, D., Webster, M., McCarthy, P., Rorden, C., Daducci, A., Alexander, D.C., Zhang, H., Dragonu, I., Matthews, P.M., Miller, K.L., Smith, S.M., 2018. Image processing and Quality Control for the first 10,000 brain imaging datasets from UK Biobank. NeuroImage 166, 400–424.

Ashburner, J., Friston, K.J., 2000. Voxel-based morphometry--the methods. NeuroImage 11, 805–821.

Bennett, C.M., Miller, M.B., 2010. How reliable are the results from functional magnetic resonance imaging. Annals of the New York Academy of Sciences 1191, 133–155.

Bossier, H., Roels, S., Seurinck, R., Banaschewski, T., Barker, G.J., Bokde, A.L.W., Quinlan, E.B., Desrivieres, S., Flor, H., Grigis, A., 2020. The empirical replicability of task-based fMRI as a function of sample size. NeuroImage 212, 116601.

Botvinik-Nezer, R., Holzmeister, F., Camerer, C.F., Dreber, A., Huber, J., Johannesson, M., Kirchler, M., Iwanir, R., Mumford, J.A., Adcock, R.A., Avesani, P., Baczkowski, B.M., Bajracharya, A., Bakst, L., Ball, S., Barilari, M., Bault, N., Beaton, D., Beitner, J., Benoit, R.G., Berkers, R.M.W.J., Bhanji, J.P., Biswal, B.B., Bobadilla-Suarez, S., Bortolini, T., Bottenhorn, K.L., Bowring, A., Braem, S., Brooks, H.R., Brudner, E.G., Calderon, C.B., Camilleri, J.A., Castrellon, J.J., Cecchetti, L., Cieslik, E.C., Cole, Z.J., Collignon, O., Cox, R.W., Cunningham, W.A., Czoschke, S., Dadi, K., Davis, C.P., Luca, A.D., Delgado, M.R., Demetriou, L., Dennison, J.B., Di, X., Dickie, E.W., Dobryakova, E., Donnat, C.L., Dukart, J., Duncan, N.W., Durnez, J., Eed, A., Eickhoff, S.B., Erhart, A., Fontanesi, L., Fricke, G.M., Fu, S., Galván, A., Gau, R., Genon, S., Glatard, T., Glerean, E., Goeman, J.J., Golowin, S.A.E., González-García, C., Gorgolewski, K.J., Grady, C.L., Green, M.A., Guassi Moreira, J.F., Guest, O., Hakimi, S., Hamilton, J.P., Hancock, R., Handjaras, G., Harry, B.B., Hawco, C., Herholz, P., Herman, G., Heunis, S., Hoffstaedter, F., Hogeveen, J., Holmes, S., Hu, C.-P., Huettel, S.A., Hughes, M.E., Iacovella, V., Iordan, A.D., Isager, P.M., Isik, A.I., Jahn, A., Johnson, M.R., Johnstone, T., Joseph, M.J.E., Juliano, A.C., Kable, J.W., Kassinopoulos, M., Koba, C., Kong, X.-Z., Koscik, T.R., Kucukboyaci, N.E., Kuhl, B.A., Kupek, S., Laird, A.R., Lamm, C., Langner, R., Lauharatanahirun, N., Lee, H., Lee, S., Leemans, A., Leo, A., Lesage, E., Li, F., Li, M.Y.C., Lim, P.C., Lintz, E.N., Liphardt, S.W., Losecaat Vermeer, A.B., Love, B.C., Mack, M.L., Malpica, N., Marins, T., Maumet, C., McDonald, K., McGuire, J.T., Melero, H., Méndez Leal, A.S., Meyer, B., Meyer, K.N., Mihai, G., Mitsis, G.D., Moll, J., Nielson, D.M., Nilsonne, G., Notter, M.P., Olivetti, E., Onicas, A.I., Papale, P., Patil, K.R., Peelle, J.E., Pérez, A., Pischedda, D., Poline, J.-B., Prystauka, Y., Ray, S., Reuter-Lorenz, P.A., Reynolds, R.C., Ricciardi, E., Rieck, J.R., Rodriguez-Thompson, A.M., Romyn, A., Salo, T., Samanez-Larkin, G.R., Sanz-Morales, E., Schlichting, M.L., Schultz, D.H., Shen, Q., Sheridan, M.A., Silvers, J.A., Skagerlund, K., Smith, A., Smith, D.V., Sokol-Hessner, P., Steinkamp, S.R., Tashjian, S.M., Thirion, B., Thorp, J.N., Tinghög, G., Tisdall, L., Tompson, S.H., Toro-Serey, C., Torre Tresols, J.J., Tozzi, L., Truong, V., Turella, L., van’t Veer, A.E., Verguts, T., Vettel, J.M., Vijayarajah, S., Vo, K., Wall, M.B., Weeda, W.D., Weis, S., White, D.J., Wisniewski, D., Xifra-Porxas, A., Yearling, E.A., Yoon, S., Yuan, R., Yuen, K.S.L., Zhang, L., Zhang, X., Zosky, J.E., Nichols, T.E., Poldrack, R.A., Schonberg, T., 2020. Variability in the analysis of a single neuroimaging dataset by many teams. Nature 582, 84–88.

Cheng, W., Palaniyappan, L., Li, M., Kendrick, K.M., Zhang, J., Luo, Q., Liu, Z., Yu, R., Deng, W., Wang, Q., Ma, X., Guo, W., Francis, S., Liddle, P., Mayer, A.R., Schumann, G., Li, T., Feng, J., 2015a. Voxel-based, brain-wide association study of aberrant functional connectivity in schizophrenia implicates thalamocortical circuitry. npj Schizophrenia 1, 15016.

Cheng, W., Rolls, E.T., Gu, H., Zhang, J., Feng, J., 2015b. Autism: reduced connectivity between cortical areas involved in face expression, theory of mind, and the sense of self. Brain 138, 1382–1393.

Efron, B., Tibshirani, R., 1997. Improvements on Cross-Validation: The 632+ Bootstrap Method. Journal of the American Statistical Association 92, 548–560.

Eklund, A., Nichols, T.E., Knutsson, H., 2016. Cluster failure: Why fMRI inferences for spatial extent have inflated false-positive rates. Proceedings of the National Academy of Sciences of the United States of America 113, 7900–7905.

Gong, W., Wan, L., Lu, W., Ma, L., Cheng, F., Cheng, W., Grunewald, S., Feng, J., 2018. Statistical testing and power analysis for brain-wide association study. Medical Image Analysis 47, 15–30.

Goodman, S., Fanelli, D., Ioannidis, J., 2016. What does research reproducibility mean? Science Translational Medicine 8, 312–341.

Greve, D.N., Fischl, B., 2009. Accurate and robust brain image alignment using boundary-based registration. NeuroImage 48, 63–72.

Lai, Y., Adam, B., Podolsky, R.H., She, J., 2007. A mixture model approach to the tests of concordance and discordance between two large-scale experiments with two-sample groups. Bioinformatics 23, 1243–1250.

Lai, Y., Eckenrode, S., She, J., 2009. A statistical framework for integrating two microarray data sets in differential expression analysis. BMC Bioinformatics 10, 1–11.

Lai, Y., Zhang, F., Nayak, T.K., Modarres, R., Lee, N.H., Mccaffrey, T.A., 2014. Concordant integrative gene set enrichment analysis of multiple large-scale two-sample expression data sets. BMC Genomics 15, 1–12.

Lai, Y., Zhang, F., Nayak, T.K., Modarres, R., Lee, N.H., Mccaffrey, T.A., 2017. An efficient concordant integrative analysis of multiple large-scale two-sample expression data sets. Bioinformatics 33, 3852–3860.

Logothetis, N.K., 2008. What we can do and what we cannot do with fMRI. Nature 453, 869–878.

Marek, K., Jennings, D., Lasch, S., Siderowf, A., Tanner, C.M., Simuni, T., Coffey, C.S., Kieburtz, K., Flagg, E., Chowdhury, S., 2011. The Parkinson Progression Marker Initiative (PPMI). Progress in Neurobiology 95, 629–635.

Mclachlan, G.J., 1987. On Bootstrapping the Likelihood Ratio Test Statistic for the Number of Components in a Normal Mixture. Applied statistics 36, 318–324.

Patel, A.X., Kundu, P., Rubinov, M., Jones, P.S., Vertes, P.E., Ersche, K.D., Suckling, J., Bullmore, E.T., 2014. A wavelet method for modeling and despiking motion artifacts from resting-state fMRI time series. NeuroImage 95, 287–304.

Poldrack, R.A., 2019. The Costs of Reproducibility. Neuron 101, 11–14.

Poldrack, R.A., Gorgolewski, K.J., 2014. Making big data open: data sharing in neuroimaging. Nature Neuroscience 17, 1510–1517.

Rolls, E.T., Joliot, M., Tzouriomazoyer, N., 2015. Implementation of a new parcellation of the orbitofrontal cortex in the automated anatomical labeling atlas. NeuroImage 122, 1–5.

Schumann, G., Loth, E., Banaschewski, T., Barbot, A., Barker, G., Büchel, C., Conrod, P.J., Dalley, J.W., Flor, H., Gallinat, J., Garavan, H., Heinz, A., Itterman, B., Lathrop, M., Mallik, C., Mann, K., Martinot, J.L., Paus, T., Poline, J.B., Robbins, T.W., Rietschel, M., Reed, L., Smolka, M., Spanagel, R., Speiser, C., Stephens, D.N., Ströhle, A., Struve, M., 2010a. The IMAGEN study: reinforcement-related behaviour in normal brain function and psychopathology. Mol Psychiatry 15, 1128–1139.

Schumann, G., Loth, E., Banaschewski, T., Barbot, A., Barker, G.J., Buchel, C., Conrod, P.J., Dalley, J.W., Flor, H., Gallinat, J., 2010b. The IMAGEN study: reinforcement-related behaviour in normal brain function and psychopathology. Molecular Psychiatry 15, 1128–1139.

Snyder, A.Z., Raichle, M.E., 2012. A brief history of the resting state: The Washington University perspective. NeuroImage 62, 902–910.

Sudlow, C., Gallacher, J., Allen, N., Beral, V., Burton, P., Danesh, J., Downey, P., Elliott, P., Green, J., Landray, M., Liu, B., Matthews, P., Ong, G., Pell, J., Silman, A., Young, A., Sprosen, T., Peakman, T., Collins, R., 2015a. UK Biobank: An Open Access Resource for Identifying the Causes of a Wide Range of Complex Diseases of Middle and Old Age. PLOS Medicine 12, e1001779.

Sudlow, C., Gallacher, J., Allen, N.E., Beral, V., Burton, P.R., Danesh, J., Downey, P., Elliott, P., Green, J., Landray, M.J., 2015b. UK biobank: an open access resource for identifying the causes of a wide range of complex diseases of middle and old age. PLOS Medicine 12.

Van Essen, D.C., Smith, S.M., Deanna, M., Behrens, T.E.J., Yacoub, E., Ugurbil, K., 2013. The WU-Minn Human Connectome Project: An Overview. NeuroImage 80, 62–79.

Van Essen, D.C., Ugurbil, K., Auerbach, E.J., Behrens, T.E.J., Bucholz, R., Chang, A., Chen, L., Corbetta, M., Curtiss, S.W., Penna, S.D., 2012. The Human Connectome Project: a data acquisition perspective. NeuroImage 62, 2222–2231.

Zhao, Y., Sampson, M.G., Wen, X., 2020. Quantify and control reproducibility in high-throughput experiments. Nature Methods 17, 1207–1213.

