## Supplementary for "Assessing Overall Reproducibility for Large-scale High-throughput MRI-based Association Studies"

### 1. Different Task for fMRI in IMAGEN

#### 1.1 The Monetary Incentive Delay Task for fMRI

Participants performed a modified version of the Monetary Incentive Delay (MID) task to examine neural responses to reward anticipation and reward outcome. The task consisted of 66 10-second trials. In each trial, participants were presented with one of three cue shapes (cue, 250 ms) denoting whether a target (white square) would subsequently appear on the left or right side of the screen and whether 0, 2 or 10 points could be won in that trial. After a variable delay (4,000-4,500 ms) of fixation on a white crosshair, participants were instructed to respond with left/right button-press as soon as the target appeared. Feedback on whether and how many points were won during the trial was presented for 1,450 ms after the response. Using a tracking algorithm, task difficulty (i.e. target duration varied between 100 and 300 ms) was individually adjusted such that each participant successfully responded on ~66% of trials. Participants had first completed a practice session outside the scanner (~5 minutes), during which they were instructed that for each 5 points won they would receive one food snack in the form of small chocolate candies.

Based on prior research suggesting reliable associations between ADHD-symptoms and fMRI BOLD responses measured during reward anticipation, the current study used the contrast 'anticipation of high-win vs anticipation of no-win'. Only successfully 'hit' trials were included here.

#### 1.2 The Emotional Reactivity fMRI Paradigm (Emotional Faces Task)

This task was adapted from 23. Participants watched 18-second blocks of either a face movie (depicting anger or neutrality) or a control stimulus. Each face movie showed black and white video clips (200-500ms) of male or female faces. Five blocks each of angry and neutral expressions were interleaved with nine blocks of the control stimulus. Each block contained eight trials of 6 face identities (3 female). The same identities were used for the angry and neutral blocks. The control stimuli were black and white concentric circles expanding and contracting at various speeds that roughly matched the contrast and motion characteristics of the face clips.

The neutral blocks contained emotional expressions that were not attributable to any particular emotion (e.g. nose twitching); however previous research has suggested that neutral stimuli are not always interpreted as such. Functional imaging studies have found significant activation of the amygdala in response to the presentation of neutral faces in healthy adult males 41, social anxiety patients and matched control participants 42, adolescents with conduct disorder problems 43 and young men with violent behavior problems 44. This suggests that neutral faces may be interpreted as emotionally ambiguous. This study focused specifically on the effects of viewing angry faces (vs control) to eliminate this ambiguity so that any significant relationships between behavior and brain could be interpreted as the consequence of viewing negative social stimuli (anger).

#### 1.3 The Stop Signal Task for fMRI

Participants performed an event-related stop signal task (SST) task designed to study neural responses to successful and unsuccessful inhibitory control 22. The task was composed of Go trials and Stop trials. During Go trials (83%; 480 trials) participants were presented with arrows pointing either to the left or to the right. During these trials, subjects were instructed to make a button response with their left or right index finger corresponding to the direction of the arrow. In the unpredictable Stop trials (17%; 80 trials), the arrows pointing left or right were followed (on average 300 ms later) by arrows pointing upwards; participants were instructed to inhibit their motor responses during these trials. A tracking algorithm changes the time interval between Go signal and Stop signal onsets according to each subject's performance on previous trials (average percentage of inhibition over previous Stop trials, recalculated after each Stop trial), resulting in 50% successful and 50% unsuccessful inhibition trials. The inter-trial interval was 1,800 ms. The tracking algorithm of the task ensured that subjects were successful on 50% of Stop trials and worked at the edge of their own inhibitory capacity.

### 2. Complete List of The IMAGEN Consortium

Tianye Jia, Alex Ing, Erin Burke Quinlan, Biondo Francesca, Tobias Banaschewski, Gareth J. Barker, Arun L. W. Bokde, Uli Bromberg, Christian Büchel, Sylvane Desrivières, Herta Flor, Hugh Garavan, Penny Gowland, Andreas Heinz, Bernd Ittermann, Jean-Luc Martinot, Frauke Nees, Dimitri Papadopoulos Orfanos, Tomáš Paus, Luise Poustka, Juliane H. Fröhner, Michael N. Smolka, Henrik Walter, Robert Whelan & Gunter Schumann.

### 3. Estimated the variance of $M^2RI$

To compute the observed information (8) within the EM framework, the objective is to find the MLE for a $p$-dimensional parameter $\theta$ in a space $\Theta$. The underlying probability model induces a density or mass function $f(x|\theta)$ on a sample space $\chi$, where $x = (x_1, \dots, x_N)^T$. Instead of observing $x \in \chi$, one observes the value of a measurable function $Y(x) = y \in Y$. The MLE $(\widetilde{\theta})$ is to be found using the data $y$.

The EM method is only attractive in situations where finding the complete data MLE and either the observed or the expected information matrix would be straightforward, but the problem based on the incomplete data $(Y)$ requires an iterative solution. Typically the complete data are from an exponential family. The algorithm operates as follows. Let

$$\lambda(x, \theta) = \log\{f(x|\theta)\}$$

$$\lambda^*(x, \theta) = \log\{f_Y(x|\theta)\} = \log\left\{\int_R f_X(x|\theta)d\mu(x)\right\}$$

where $R = x : y(x) = y$, and $\mu(x)$ is a dominating measure.

To see how to compute the observed information in the EM, let $S(x, \theta)$ and $S^*(y, \theta)$ be the gradient vectors of $\lambda$ and $\lambda^*$ respectively and $B(x, \theta)$ and $B^*(y, \theta)$ be the negatives of the associated second derivative matrices. Then by straightforward differentiation:

$$S^*(y, \theta) = E_\theta[S(X, \theta)|X \in R]$$

$$S^*(y, \widetilde{\theta}) = 0$$

$$I_Y(\theta) = E_\theta\{B(X, \theta)|X \in R\} - E_\theta\left\{S(X, \theta)S^T(X, \theta)|X \in R\right\} + S^*(X, \theta)S^{*T}(X, \theta)$$

Assume data $Z = \left\{[Z_1^{(1)}, Z_1^{(2)}], \dots, [Z_N^{(1)}, Z_N^{(2)}]\right\}$ are known to be i.i.d. from our model, this can be considered a missing data problem by letting $X = (Z, W)$, where

$$W = W_{ij}^k = \begin{cases} 1, & k\text{th data point from } (i, j) \text{ component} \\ 0, & \text{otherwise} \end{cases}$$

Then we have

$$\lambda(X|\Theta) = \lambda(Z, W|\Theta) = \log \prod_{k=1}^N \prod_{i=0}^2 \prod_{j=0}^2 \{\pi_{ij}\phi_{\mu_i, \sigma_i^2}[z_n^{(1)}]\phi_{\nu_j, \tau_j^2}[z_n^{(2)}]\}^{w_{ij}^n}$$

$$= \sum_{k=1}^{N}\{\sum_{i=0}^{2}\sum_{j=0}^{2} W_{ij}^{k} \log \pi_{ij} + \sum_{i=0}^{2}\sum_{j=0}^{2} W_{ij}^{k}[\log \phi_{\mu_i,\sigma_i^2}[z_n^{(1)}] + \log \phi_{\nu_j,\tau_j^2}[z_n^{(2)}]]\}$$

Let

$$\pi_{00} = 1 - \sum_{i+j\neq 0} \pi_{ij}$$

Where $\Theta = \{\pi_{ij}, \mu_1, \mu_2, \nu_1, \nu_2, \sigma_1^2, \sigma_2^2, \tau_1^2, \tau_2^2\}, i \in \{0,1,2\}, j \in \{0,1,2\}$ and $i + j \neq 0$.

Separate the component $i + j = 0$, then we have

$$\lambda(Z, W|\Theta) = \sum_{k=1}^{N}\{\sum_{i+j\neq 0} W_{ij}^{k} \log \pi_{ij} + (1 - \sum_{i+j\neq 0} W_{ij}^{k}) \log(1 - \sum_{i+j\neq 0} \pi_{ij})$$

$$+ \sum_{i+j\neq 0} W_{ij}^{k}[-\frac{1}{2}\log(2\pi\sigma_i^2) - \frac{1}{2}\log(2\pi\tau_i^2) - \frac{(Z_k^{(1)} - \mu_i)^2}{2\sigma_i^2} - \frac{(Z_k^{(2)} - \mu_j)^2}{2\tau_j^2}]\}$$

$$+(1 - \sum_{i+j\neq 0} W_{ij}^{k})[-\log(2\pi) - \frac{(Z_k^{(1)})^2 + (Z_k^{(2)})^2}{2}]$$

**Part I** $E_\theta\{B(X,\theta)|X \in R\}$

1)

$$B_{(i,j)=(h,l)}(\pi_{ij}, \pi_{hl}) = -\frac{\partial^2}{\partial \pi_{ij}^2} = \sum_{k=1}^{N}\{\frac{W_{ij}^{k}}{\pi_{ij}^2} + \frac{W_{00}^{k}}{\pi_{00}^2}\}$$

$$E_\theta\{B_{(i,j)=(h,l)}(\pi_{ij}, \pi_{hl})\} = \sum_{k=1}^{N}\{\frac{E[W_{ij}^{k}]}{\pi_{ij}^2} + \frac{E[W_{00}^{k}]}{\pi_{00}^2}\}$$

Where $i, h \in \{0,1,2\}, j, l \in \{0,1,2\}$, $h + l \neq 0$ and $i + j \neq 0$.

2)

$$B_{(i,j)\neq(h,l)}(\pi_{ij}, \pi_{hl}) = -\frac{\partial^2}{\partial \pi_{ij}\partial \pi_{hl}} = \sum_{k=1}^{N}\{\frac{W_{00}^{k}}{\pi_{00}^2}\}$$

$$E_\theta\{B_{(i,j)\neq(h,l)}(\pi_{ij}, \pi_{hl})\} = \sum_{k=1}^{N}\{\frac{E[W_{00}^{k}]}{\pi_{00}^2}\}$$

Where $i, h \in \{0,1,2\}, j, l \in \{0,1,2\}$, $h + l \neq 0$ and $i + j \neq 0$.

3)

$$B_{i=h}(\mu_i, \mu_h) = -\frac{\partial^2}{\partial \mu_i^2} = \sum_{k=1}^{N}\sum_{j=0}^{2}\{\frac{W_{ij}^{k}}{\sigma_i^2}\}$$

$$E_\theta\{B_{i=h}(\mu_i,\mu_h)\} = \sum_{k=1}^{N}\sum_{j=0}^{2}\{\frac{E[W_{ij}^k]}{\sigma_i^2}\}$$

Where $i, h \in \{1,2\}, j, l \in \{0,1,2\}$.

4)

$$B_{j=l}(\nu_j,\nu_l) = -\frac{\partial^2}{\partial \nu_j^2} = \sum_{k=1}^{N}\sum_{i=0}^{2}\{\frac{W_{ij}^k}{\tau_j^2}\}$$

$$E_\theta\{B_{j=l}(\mu_j,\mu_l)\} = \sum_{k=1}^{N}\sum_{i=0}^{2}\{\frac{E[W_{ij}^k]}{\tau_j^2}\}$$

Where $i, h \in \{0,1,2\}, j, l \in \{1,2\}$.

5)

$$B_{i=h}(\sigma_i^2,\sigma_h^2) = -\frac{\partial^2}{\partial(\sigma_i^2)^2} = \sum_{k=1}^{N}\sum_{j=0}^{2}\{W_{ij}^k[\frac{(Z_k^{(1)}-\mu_i)^2}{(\sigma_i^2)^3} - \frac{1}{2(\sigma_i^2)^2}]\}$$

$$E_\theta\{B_{i=h}(\sigma_i^2,\sigma_h^2)\} = \sum_{k=1}^{N}\sum_{j=0}^{2}\{E[W_{ij}^k][\frac{(Z_k^{(1)}-\mu_i)^2}{(\sigma_i^2)^3} - \frac{1}{2(\sigma_i^2)^2}]\}$$

Where $i, h \in \{1,2\}, j, l \in \{0,1,2\}$.

6)

$$B_{j=l}(\tau_j^2,\tau_l^2) = -\frac{\partial^2}{\partial(\tau_j^2)^2} = \sum_{k=1}^{N}\sum_{i=0}^{2}\{W_{ij}^k[\frac{(Z_k^{(2)}-\nu_j)^2}{(\tau_j^2)^3} - \frac{1}{2(\tau_j^2)^2}]\}$$

$$E_\theta\{B_{j=l}(\tau_j^2,\tau_l^2)\} = \sum_{k=1}^{N}\sum_{i=0}^{2}\{E[W_{ij}^k][\frac{(Z_k^{(2)}-\nu_j)^2}{(\tau_j^2)^3} - \frac{1}{2(\tau_j^2)^2}]\}$$

Where $i, h \in \{0,1,2\}, j, l \in \{1,2\}$.

7)

$$B_{i=h}(\mu_i,\sigma_h^2) = -\frac{\partial^2}{\partial\mu_i\partial\sigma_i^2} = \sum_{k=1}^{N}\sum_{j=0}^{2}\{W_{ij}^k[\frac{Z_k^{(1)}-\mu_i}{(\sigma_i^2)^2}]\}$$

$$E_\theta\{B_{i=h}(\mu_i,\sigma_h^2)\} = \sum_{k=1}^{N}\sum_{j=0}^{2}\{E[W_{ij}^k][\frac{Z_k^{(1)}-\mu_i}{(\sigma_i^2)^2}]\}$$

Where $i, h \in \{1,2\}, j, l \in \{0,1,2\}$.

8)

$$B_{j=l}(\nu_j,\tau_l^2) = -\frac{\partial^2}{\partial\nu_j\partial\tau_j^2} = \sum_{k=1}^{N}\sum_{j=0}^{2}\{W_{ij}^k[\frac{Z_k^{(2)}-\nu_j}{(\tau_j^2)^2}]\}$$

$$E_\theta\{B_{j=l}(\nu_j,\tau_l^2)\} = \sum_{k=1}^{N}\sum_{i=0}^{2}\{E[W_{ij}^k][\frac{Z_k^{(2)}-\nu_j}{(\tau_j^2)^2}]\}$$

Where $i,h \in \{0,1,2\}, j,l \in \{1,2\}$.

9) Otherwise equal to 0.

**Part *II* $\boldsymbol{E_\theta\left\{S(X,\theta)S^T(X,\theta)|X \in R\right\}}$**

1)

$$S(\pi_{ij})S(\pi_{ij}) = \frac{\partial}{\partial\pi_{ij}} \times \frac{\partial}{\partial\pi_{ij}} =$$

$$[\sum_{k_1=1}^{N}(\frac{W_{ij}^{k_1}}{\pi_{ij}} - \frac{W_{00}^{k_1}}{\pi_{00}})][\sum_{k_2=1}^{N}(\frac{W_{ij}^{k_2}}{\pi_{ij}} - \frac{W_{00}^{k_2}}{\pi_{00}})]$$

Where $i,h \in \{0,1,2\}, j,l \in \{0,1,2\}, i+j \neq 0$ and $h+l \neq 0$.

$$E[S(\pi_{ij})S(\pi_{ij})] = E[\frac{\partial}{\partial\pi_{ij}} \times \frac{\partial}{\partial\pi_{ij}}] =$$

$$E\{[\sum_{k_1=1}^{N}(\frac{W_{ij}^{k_1}}{\pi_{ij}} - \frac{W_{00}^{k_1}}{\pi_{00}})][\sum_{k_2=1}^{N}(\frac{W_{ij}^{k_2}}{\pi_{ij}} - \frac{W_{00}^{k_2}}{\pi_{00}})]\}$$

$$= \sum_{k_1\neq k_2}[\frac{E[W_{ij}^{k_1}W_{ij}^{k_2}]}{\pi_{ij}^2} - \frac{E[W_{ij}^{k_1}W_{00}^{k_2}]}{\pi_{ij}\pi_{00}} - \frac{E[W_{00}^{k_1}W_{ij}^{k_2}]}{\pi_{00}\pi_{ij}} + \frac{E[W_{00}^{k_1}W_{00}^{k_2}]}{\pi_{00}^2}]$$

$$+ \sum_{k_1=k_2=k}[\frac{E[W_{ij}^{k}W_{ij}^{k}]}{\pi_{ij}^2} - \frac{2E[W_{ij}^{k}W_{00}^{k}]}{\pi_{ij}\pi_{00}} + \frac{E[W_{00}^{k}W_{00}^{k}]}{\pi_{00}^2}]$$

As $W_{ij}^k, W_{00}^k \in \{0,1\}$, we have $E[W_{ij}^kW_{ij}^k] = E[W_{ij}^k]$ and $E[W_{00}^kW_{00}^k] = E[W_{00}^k]$. What's more, $E[W_{ij}^kW_{00}^k] = 0$ because $W_{ij}^k, W_{00}^k$ can not equal to 1 at the same time. Then we have

$$E[S(\pi_{ij})S(\pi_{ij})]$$

$$= \sum_{k_1\neq k_2}[\frac{E[W_{ij}^{k_1}]E[W_{ij}^{k_2}]}{\pi_{ij}^2} - \frac{E[W_{ij}^{k_1}]E[W_{00}^{k_2}]}{\pi_{ij}\pi_{00}} - \frac{E[W_{00}^{k_1}]E[W_{ij}^{k_2}]}{\pi_{00}\pi_{ij}} + \frac{E[W_{00}^{k_1}]E[W_{00}^{k_2}]}{\pi_{00}^2}]$$

$$+ \sum_{k_1=k_2=k}[\frac{E[W_{ij}^{k}]}{\pi_{ij}^2} + \frac{E[W_{00}^{k}]}{\pi_{00}^2}]$$

2)

$$S(\pi_{ij})S(\pi_{hl}) = \frac{\partial}{\partial\pi_{ij}} \times \frac{\partial}{\partial\pi_{hl}} =$$

$$[\sum_{k_1=1}^{N}(\frac{W_{ij}^{k_1}}{\pi_{ij}}-\frac{W_{00}^{k_1}}{\pi_{00}})][\sum_{k_2=1}^{N}(\frac{W_{hl}^{k_2}}{\pi_{hl}}-\frac{W_{00}^{k_2}}{\pi_{00}})]$$

Where $i,h \in \{0,1,2\}, j,l \in \{0,1,2\}, i+j \neq 0, h+l \neq 0$ and $(i,j) \neq (h,l)$.

$$E[S(\pi_{ij})S(\pi_{hl})] = E[\frac{\partial}{\partial \pi_{ij}} \times \frac{\partial}{\partial \pi_{hl}}] =$$

$$E\{[\sum_{k_1=1}^{N}(\frac{W_{ij}^{k_1}}{\pi_{ij}}-\frac{W_{00}^{k_1}}{\pi_{00}})][\sum_{k_2=1}^{N}(\frac{W_{hl}^{k_2}}{\pi_{hl}}-\frac{W_{00}^{k_2}}{\pi_{00}})]\}$$

$$= \sum_{k_1 \neq k_2}[\frac{E[W_{ij}^{k_1}W_{hl}^{k_2}]}{\pi_{ij}\pi_{hl}} - \frac{E[W_{ij}^{k_1}W_{00}^{k_2}]}{\pi_{ij}\pi_{00}} - \frac{E[W_{00}^{k_1}W_{hl}^{k_2}]}{\pi_{00}\pi_{hl}} + \frac{E[W_{00}^{k_1}W_{00}^{k_2}]}{\pi_{00}^2}]$$

$$+ \sum_{k_1=k_2=k}[\frac{E[W_{ij}^{k}W_{hl}^{k}]}{\pi_{ij}\pi_{hl}} - \frac{E[W_{ij}^{k}W_{00}^{k}]}{\pi_{ij}\pi_{00}} - \frac{E[W_{00}^{k}W_{hl}^{k}]}{\pi_{00}\pi_{hl}} + \frac{E[W_{00}^{k}W_{00}^{k}]}{\pi_{00}^2}]$$

As $W_{ij}^k, W_{hl}^k, W_{00}^k \in \{0,1\}$, we have $E[W_{00}^k W_{00}^k] = E[W_{00}^k]$. What's more, $E[W_{ij}^k W_{hl}^k] = 0$, $E[W_{00}^k W_{hl}^k] = 0$ and $E[W_{ij}^k W_{00}^k] = 0$ because $W_{ij}^k, W_{hl}^k$ and $W_{00}^k$ can not equal to 1 at the same time. Then we have

$$E[S(\pi_{ij})S(\pi_{hl})]$$

$$= \sum_{k_1 \neq k_2}[\frac{E[W_{ij}^{k_1}]E[W_{hl}^{k_2}]}{\pi_{ij}\pi_{hl}} - \frac{E[W_{ij}^{k_1}]E[W_{00}^{k_2}]}{\pi_{ij}\pi_{00}} - \frac{E[W_{00}^{k_1}]E[W_{hl}^{k_2}]}{\pi_{00}\pi_{hl}} + \frac{E[W_{00}^{k_1}]E[W_{00}^{k_2}]}{\pi_{00}^2}]$$

$$+ \sum_{k_1=k_2=k}[\frac{E[W_{00}^k]}{\pi_{00}^2}]$$

3)

$$S(\mu_i)S(\mu_i) = \frac{\partial}{\partial \mu_i} \times \frac{\partial}{\partial \mu_i} =$$

$$[\sum_{k_1=1}^{N}\sum_{j=0}^{2} W_{ij}^{k_1}\frac{Z_{k_1}^{(1)}-\mu_i}{\sigma_i^2}][\sum_{k_2=1}^{N}\sum_{l=0}^{2} W_{il}^{k_2}\frac{Z_{k_2}^{(1)}-\mu_i}{\sigma_i^2}]$$

Where $i \in \{1,2\}$ and $j,l \in \{0,1,2\}$.

$$E[S(\mu_i)S(\mu_i)] = E[\frac{\partial}{\partial \mu_i} \times \frac{\partial}{\partial \mu_i}] =$$

$$E\{[\sum_{k_1=1}^{N}\sum_{j=0}^{2} W_{ij}^{k_1}\frac{Z_{k_1}^{(1)}-\mu_i}{\sigma_i^2}][\sum_{k_2=1}^{N}\sum_{l=0}^{2} W_{il}^{k_2}\frac{Z_{k_2}^{(1)}-\mu_i}{\sigma_i^2}]\}$$

$$= \sum_{k_1 \neq k_2}[\sum_{j=0}^{2}\sum_{l=0}^{2} E[W_{ij}^{k_1}W_{il}^{k_2}]\frac{Z_{k_1}^{(1)}-\mu_i}{\sigma_i^2}\frac{Z_{k_2}^{(1)}-\mu_i}{\sigma_i^2}]$$

$$+\sum_{k_1=k_2=k}[\sum_{j=0}^{2}\sum_{l=0}^{2}E[W_{ij}^{k}W_{il}^{k}](\frac{Z_k^{(1)}-\mu_i}{\sigma_i^2})^2]$$

As $W_{ij}^k \in \{0,1\}$, when $j=l$ we have $E[W_{ij}^kW_{il}^k]=E[W_{ij}^kW_{ij}^k]=E[W_{ij}^k]$. What's more, when $j \neq l$ we have $E[W_{ij}^kW_{il}^k]=0$ because $W_{ij}^k, W_{il}^k \in \{0,1\}$ and $W_{ij}^k, W_{il}^k$ not equal to 1 at the same time. Then we have

$$E[S(\mu_i)S(\mu_i)]$$

$$=\sum_{k_1\neq k_2}[\sum_{j=0}^{2}\sum_{l=0}^{2}E[W_{ij}^{k_1}]E[W_{il}^{k_2}]\frac{Z_{k_1}^{(1)}-\mu_i}{\sigma_i^2}\frac{Z_{k_2}^{(1)}-\mu_i}{\sigma_i^2}]$$

$$+\sum_{k_1=k_2=k}[\sum_{j=0}^{2}E[W_{ij}^{k}](\frac{Z_k^{(1)}-\mu_i}{\sigma_i^2})^2]$$

4)

$$S(\mu_i)S(\mu_h)=\frac{\partial}{\partial\mu_i}\times\frac{\partial}{\partial\mu_h}=$$

$$[\sum_{k_1=1}^{N}\sum_{j=0}^{2}W_{ij}^{k_1}\frac{Z_{k_1}^{(1)}-\mu_i}{\sigma_i^2}][\sum_{k_2=1}^{N}\sum_{l=0}^{2}W_{hl}^{k_2}\frac{Z_{k_2}^{(1)}-\mu_h}{\sigma_h^2}]$$

Where $i,h\in\{1,2\}$, $j,l\in\{0,1,2\}$ and $i\neq h$.

$$E[S(\mu_i)S(\mu_h)]=E[\frac{\partial}{\partial\mu_i}\times\frac{\partial}{\partial\mu_h}]=$$

$$E\{[\sum_{k_1=1}^{N}\sum_{j=0}^{2}W_{ij}^{k_1}\frac{Z_{k_1}^{(1)}-\mu_i}{\sigma_i^2}][\sum_{k_2=1}^{N}\sum_{l=0}^{2}W_{hl}^{k_2}\frac{Z_{k_2}^{(1)}-\mu_h}{\sigma_h^2}]\}$$

$$=\sum_{k_1\neq k_2}[\sum_{j=0}^{2}\sum_{l=0}^{2}E[W_{ij}^{k_1}W_{hl}^{k_2}]\frac{Z_{k_1}^{(1)}-\mu_i}{\sigma_i^2}\frac{Z_{k_2}^{(1)}-\mu_h}{\sigma_h^2}]$$

$$+\sum_{k_1=k_2=k}[\sum_{j=0}^{2}\sum_{l=0}^{2}E[W_{ij}^{k}W_{hl}^{k}]\frac{Z_k^{(1)}-\mu_i}{\sigma_i^2}\frac{Z_k^{(1)}-\mu_h}{\sigma_h^2}]$$

As $W_{ij}^k, W_{hl}^k \in \{0,1\}$, $E[W_{ij}^kW_{hl}^k]=0$ because $W_{ij}^k, W_{hl}^k$ can not be 1 at the same time. Then we have

$$E[S(\mu_i)S(\mu_h)]$$

$$=\sum_{k_1\neq k_2}[\sum_{j=0}^{2}\sum_{l=0}^{2}E[W_{ij}^{k_1}]E[W_{hl}^{k_2}]\frac{Z_{k_1}^{(1)}-\mu_i}{\sigma_i^2}\frac{Z_{k_2}^{(1)}-\mu_h}{\sigma_h^2}]$$

5)

$$S(\nu_j)S(\nu_j) = \frac{\partial}{\partial \nu_j} \times \frac{\partial}{\partial \nu_j} =$$

$$[\sum_{k_1=1}^{N}\sum_{i=0}^{2} W_{ij}^{k_1} \frac{Z_{k_1}^{(2)} - \nu_j}{\tau_j^2}][\sum_{k_2=1}^{N}\sum_{h=0}^{2} W_{hj}^{k_2} \frac{Z_{k_2}^{(2)} - \nu_j}{\tau_j^2}]$$

Where $i, h \in \{0, 1, 2\}$ and $j \in \{1, 2\}$.

$$E[S(\nu_j)S(\nu_j)] = E[\frac{\partial}{\partial \nu_j} \times \frac{\partial}{\partial \nu_j}] =$$

$$E\{[\sum_{k_1=1}^{N}\sum_{i=0}^{2} W_{ij}^{k_1} \frac{Z_{k_1}^{(2)} - \nu_j}{\tau_j^2}][\sum_{k_2=1}^{N}\sum_{h=0}^{2} W_{hj}^{k_2} \frac{Z_{k_2}^{(2)} - \nu_j}{\tau_j^2}]\}$$

$$= \sum_{k_1 \neq k_2} [\sum_{i=0}^{2}\sum_{h=0}^{2} E[W_{ij}^{k_1} W_{hj}^{k_2}] \frac{Z_{k_1}^{(2)} - \nu_j}{\tau_j^2} \frac{Z_{k_2}^{(2)} - \nu_j}{\tau_j^2}]$$

$$+ \sum_{k_1=k_2=k} [\sum_{i=0}^{2}\sum_{h=0}^{2} E[W_{ij}^{k} W_{hj}^{k}] (\frac{Z_{k}^{(2)} - \nu_j}{\tau_j^2})^2]$$

As $W_{ij}^k \in \{0, 1\}$, when $i = h$ we have $E[W_{ij}^k W_{hj}^k] = E[W_{ij}^k W_{ij}^k] = E[W_{ij}^k]$. What's more, when $j \neq l$ we have $E[W_{ij}^k W_{hj}^k] = 0$ because $W_{ij}^k, W_{hj}^k \in \{0, 1\}$ and $W_{ij}^k, W_{hj}^k$ not equal to 1 at the same time. Then we have

$$E[S(\nu_i)S(\nu_j)]$$

$$= \sum_{k_1 \neq k_2} [\sum_{i=0}^{2}\sum_{h=0}^{2} E[W_{ij}^{k_1}] E[W_{hj}^{k_2}] \frac{Z_{k_1}^{(2)} - \nu_j}{\tau_j^2} \frac{Z_{k_2}^{(2)} - \nu_j}{\tau_j^2}]$$

$$+ \sum_{k_1=k_2=k} [\sum_{i=0}^{2} E[W_{ij}^{k}] (\frac{Z_{k}^{(2)} - \nu_j}{\tau_j^2})^2]$$

6)

$$S(\nu_j)S(\nu_l) = \frac{\partial}{\partial \nu_j} \times \frac{\partial}{\partial \nu_l} =$$

$$[\sum_{k_1=1}^{N}\sum_{i=0}^{2} W_{ij}^{k_1} \frac{Z_{k_1}^{(2)} - \nu_j}{\tau_j^2}][\sum_{k_2=1}^{N}\sum_{h=0}^{2} W_{hl}^{k_2} \frac{Z_{k_2}^{(2)} - \nu_l}{\tau_l^2}]$$

Where $i, h \in \{0, 1, 2\}$, $j, l \in \{1, 2\}$ and $(i, j) \neq (h, l)$.

$$E[S(\nu_j)S(\nu_l)] = E[\frac{\partial}{\partial \nu_j} \times \frac{\partial}{\partial \nu_l}] =$$

$$E\{[\sum_{k_1=1}^{N}\sum_{i=0}^{2} W_{ij}^{k_1}\frac{Z_{k_1}^{(2)}-\nu_j}{\tau_j^2}][\sum_{k_2=1}^{N}\sum_{h=0}^{2} W_{hl}^{k_2}\frac{Z_{k_2}^{(2)}-\nu_l}{\tau_l^2}]\}$$

$$= \sum_{k_1\neq k_2}[\sum_{i=0}^{2}\sum_{h=0}^{2} E[W_{ij}^{k_1}W_{hl}^{k_2}]\frac{Z_{k_1}^{(2)}-\nu_j}{\tau_j^2}\frac{Z_{k_2}^{(2)}-\nu_l}{\tau_l^2}]$$

$$+ \sum_{k_1=k_2=k}[\sum_{i=0}^{2}\sum_{h=0}^{2} E[W_{ij}^{k}W_{hl}^{k}]\frac{Z_{k}^{(2)}-\nu_j}{\tau_j^2}\frac{Z_{k}^{(2)}-\nu_l}{\tau_l^2}]$$

As $W_{ij}^k, W_{hl}^k \in \{0,1\}$, $E[W_{ij}^k W_{hl}^k] = 0$ because $W_{ij}^k, W_{hl}^k$ can not be 1 at the same time. Then we have

$$E[S(\nu_j)S(\nu_l)]$$

$$= \sum_{k_1\neq k_2}[\sum_{i=0}^{2}\sum_{h=0}^{2} E[W_{ij}^{k_1}]E[W_{hl}^{k_2}]\frac{Z_{k_1}^{(2)}-\nu_j}{\tau_j^2}\frac{Z_{k_2}^{(2)}-\nu_l}{\tau_l^2}]$$

7)

$$S(\mu_i)S(\nu_l) = \frac{\partial}{\partial \mu_i}\times\frac{\partial}{\partial \nu_l} =$$

$$[\sum_{k_1=1}^{N}\sum_{j=0}^{2} W_{ij}^{k_1}\frac{Z_{k_1}^{(1)}-\mu_i}{\sigma_i^2}][\sum_{k_2=1}^{N}\sum_{h=0}^{2} W_{hl}^{k_2}\frac{Z_{k_2}^{(2)}-\nu_l}{\tau_l^2}]$$

Where $j, h \in \{0,1,2\}$, $i, l \in \{1,2\}$.

$$E[S(\mu_i)S(\nu_l)] = E[\frac{\partial}{\partial \mu_i}\times\frac{\partial}{\partial \nu_l}] =$$

$$E\{[\sum_{k_1=1}^{N}\sum_{j=0}^{2} W_{ij}^{k_1}\frac{Z_{k_1}^{(1)}-\mu_i}{\sigma_i^2}][\sum_{k_2=1}^{N}\sum_{h=0}^{2} W_{hl}^{k_2}\frac{Z_{k_2}^{(2)}-\nu_l}{\tau_l^2}]\}$$

$$= \sum_{k_1\neq k_2}[\sum_{j=0}^{2}\sum_{h=0}^{2} E[W_{ij}^{k_1}W_{hl}^{k_2}]\frac{Z_{k_1}^{(1)}-\mu_i}{\sigma_i^2}\frac{Z_{k_2}^{(2)}-\nu_l}{\tau_l^2}]$$

$$+ \sum_{k_1=k_2=k}[\sum_{j=0}^{2}\sum_{h=0}^{2} E[W_{ij}^{k}W_{hl}^{k}]\frac{Z_{k}^{(1)}-\mu_i}{\sigma_i^2}\frac{Z_{k}^{(2)}-\nu_l}{\tau_l^2}]$$

$E[W_{ij}^k W_{hl}^k] = E[W_{il}^k]$ because $W_{ij}^k, W_{hl}^k \in \{0,1\}$ can not be 1 at the same time unless $(i,j) = (h,l) = (i,l)$. Then we have

$$E[S(\mu_i)S(\nu_l)]$$

$$= \sum_{k_1\neq k_2}[\sum_{j=0}^{2}\sum_{h=0}^{2} E[W_{ij}^{k_1}]E[W_{hl}^{k_2}]\frac{Z_{k_1}^{(1)}-\mu_i}{\sigma_i^2}\frac{Z_{k_2}^{(2)}-\nu_l}{\tau_l^2}]$$

$$+ \sum_{k_1=k_2=k}[E[W_{il}^{k}]\frac{Z_{k}^{(1)}-\mu_i}{\sigma_i^2}\frac{Z_{k}^{(2)}-\nu_l}{\tau_l^2}]$$

8)

$$S(\sigma_i^2)S(\sigma_i^2) = \frac{\partial}{\partial \sigma_i^2} \times \frac{\partial}{\partial \sigma_i^2} =$$

$$[\sum_{k_1=1}^{N}\sum_{j=0}^{2} W_{ij}^{k_1}[\frac{(Z_{k_1}^{(1)}-\mu_i)^2}{2(\sigma_i^2)^2} - \frac{1}{2\sigma_i^2}]][\sum_{k_2=1}^{N}\sum_{l=0}^{2} W_{il}^{k_2}[\frac{(Z_{k_2}^{(1)}-\mu_i)^2}{2(\sigma_i^2)^2} - \frac{1}{2\sigma_i^2}]]$$

Where $i \in \{1,2\}$ and $j,l \in \{0,1,2\}$.

$$E[S(\sigma_i^2)S(\sigma_i^2)] = E[\frac{\partial}{\partial \sigma_i^2} \times \frac{\partial}{\partial \sigma_i^2}]$$

$$= E\{[\sum_{k_1=1}^{N}\sum_{j=0}^{2} W_{ij}^{k_1}[\frac{(Z_{k_1}^{(1)}-\mu_i)^2}{2(\sigma_i^2)^2} - \frac{1}{2\sigma_i^2}]][\sum_{k_2=1}^{N}\sum_{l=0}^{2} W_{il}^{k_2}[\frac{(Z_{k_2}^{(1)}-\mu_i)^2}{2(\sigma_i^2)^2} - \frac{1}{2\sigma_i^2}]]\}$$

$$= \sum_{k_1 \neq k_2}[\sum_{j=0}^{2}\sum_{l=0}^{2} E[W_{ij}^{k_1}W_{il}^{k_2}][\frac{(Z_{k_1}^{(1)}-\mu_i)^2}{2(\sigma_i^2)^2} - \frac{1}{2\sigma_i^2}][\frac{(Z_{k_2}^{(1)}-\mu_i)^2}{2(\sigma_i^2)^2} - \frac{1}{2\sigma_i^2}]]$$

$$+ \sum_{k_1=k_2=k}[\sum_{j=0}^{2}\sum_{l=0}^{2} E[W_{ij}^{k}W_{il}^{k}](\frac{(Z_{k}^{(1)}-\mu_i)^2}{2(\sigma_i^2)^2} - \frac{1}{2\sigma_i^2})^2]$$

As $W_{ij}^k \in \{0,1\}$, when $j = l$ we have $E[W_{ij}^k W_{il}^k] = E[W_{ij}^k W_{ij}^k] = E[W_{ij}^k]$. What's more, when $j \neq l$ we have $E[W_{ij}^k W_{il}^k] = 0$ because $W_{ij}^k, W_{il}^k \in \{0,1\}$ and $W_{ij}^k, W_{il}^k$ not equal to 1 at the same time. Then we have

$$E[S(\sigma_i^2)S(\sigma_i^2)]$$

$$= \sum_{k_1 \neq k_2}[\sum_{j=0}^{2}\sum_{l=0}^{2} E[W_{ij}^{k_1}]E[W_{il}^{k_2}][\frac{(Z_{k_1}^{(1)}-\mu_i)^2}{2(\sigma_i^2)^2} - \frac{1}{2\sigma_i^2}][\frac{(Z_{k_2}^{(1)}-\mu_i)^2}{2(\sigma_i^2)^2} - \frac{1}{2\sigma_i^2}]]$$

$$+ \sum_{k_1=k_2=k}[\sum_{j=0}^{2} E[W_{ij}^{k}](\frac{(Z_{k}^{(1)}-\mu_i)^2}{2(\sigma_i^2)^2} - \frac{1}{2\sigma_i^2})^2]$$

9)

$$S(\sigma_i^2)S(\sigma_h^2) = \frac{\partial}{\partial \sigma_i^2} \times \frac{\partial}{\partial \sigma_h^2} =$$

$$[\sum_{k_1=1}^{N}\sum_{j=0}^{2} W_{ij}^{k_1}[\frac{(Z_{k_1}^{(1)}-\mu_i)^2}{2(\sigma_i^2)^2} - \frac{1}{2\sigma_i^2}]][\sum_{k_2=1}^{N}\sum_{l=0}^{2} W_{hl}^{k_2}[\frac{(Z_{k_2}^{(1)}-\mu_h)^2}{2(\sigma_h^2)^2} - \frac{1}{2\sigma_h^2}]]$$

Where $i,h \in \{1,2\}$, $j,l \in \{0,1,2\}$ and $i \neq h$.

$$E[S(\sigma_i^2)S(\sigma_h^2)] = E[\frac{\partial}{\partial \sigma_i^2} \times \frac{\partial}{\partial \sigma_h^2}]$$

$$= E\{[\sum_{k_1=1}^{N}\sum_{j=0}^{2} W_{ij}^{k_1}[\frac{(Z_{k_1}^{(1)}-\mu_i)^2}{2(\sigma_i^2)^2} - \frac{1}{2\sigma_i^2}]][\sum_{k_2=1}^{N}\sum_{l=0}^{2} W_{hl}^{k_2}[\frac{(Z_{k_2}^{(1)}-\mu_h)^2}{2(\sigma_h^2)^2} - \frac{1}{2\sigma_h^2}]]\}$$

$$= \sum_{k_1 \neq k_2}[\sum_{j=0}^{2}\sum_{l=0}^{2} E[W_{ij}^{k_1}W_{hl}^{k_2}][\frac{(Z_{k_1}^{(1)}-\mu_i)^2}{2(\sigma_i^2)^2} - \frac{1}{2\sigma_i^2}][\frac{(Z_{k_2}^{(1)}-\mu_h)^2}{2(\sigma_h^2)^2} - \frac{1}{2\sigma_h^2}]]$$

$$+ \sum_{k_1=k_2=k}[\sum_{j=0}^{2}\sum_{l=0}^{2} E[W_{ij}^{k}W_{hl}^{k}][\frac{(Z_{k}^{(1)}-\mu_i)^2}{2(\sigma_i^2)^2} - \frac{1}{2\sigma_i^2}][\frac{(Z_{k}^{(1)}-\mu_h)^2}{2(\sigma_h^2)^2} - \frac{1}{2\sigma_h^2}]]$$

As $W_{ij}^k, W_{hl}^k \in \{0,1\}$, $E[W_{ij}^k W_{hl}^k] = 0$ because $W_{ij}^k, W_{hl}^k$ can not be 1 at the same time. Then we have

$$E[S(\sigma_i^2)S(\sigma_h^2)]$$

$$= \sum_{k_1 \neq k_2}[\sum_{j=0}^{2}\sum_{l=0}^{2} E[W_{ij}^{k_1}]E[W_{hl}^{k_2}][\frac{(Z_{k_1}^{(1)}-\mu_i)^2}{2(\sigma_i^2)^2} - \frac{1}{2\sigma_i^2}][\frac{(Z_{k_2}^{(1)}-\mu_h)^2}{2(\sigma_h^2)^2} - \frac{1}{2\sigma_h^2}]]$$

10)

$$S(\tau_j^2)S(\tau_j^2) = \frac{\partial}{\partial \tau_j^2} \times \frac{\partial}{\partial \tau_j^2} =$$

$$[\sum_{k_1=1}^{N}\sum_{i=0}^{2} W_{ij}^{k_1}[\frac{(Z_{k_1}^{(2)}-\nu_j)^2}{2(\tau_j^2)^2} - \frac{1}{2\tau_j^2}]][\sum_{k_2=1}^{N}\sum_{h=0}^{2} W_{hj}^{k_2}[\frac{(Z_{k_2}^{(2)}-\nu_j)^2}{2(\tau_j^2)^2} - \frac{1}{2\tau_j^2}]]$$

Where $i, h \in \{0,1,2\}$ and $j \in \{1,2\}$.

$$E[S(\tau_j^2)S(\tau_j^2)] = E[\frac{\partial}{\partial \tau_j^2} \times \frac{\partial}{\partial \tau_j^2}]$$

$$= E\{[\sum_{k_1=1}^{N}\sum_{i=0}^{2} W_{ij}^{k_1}[\frac{(Z_{k_1}^{(2)}-\nu_j)^2}{2(\tau_j^2)^2} - \frac{1}{2\tau_j^2}]][\sum_{k_2=1}^{N}\sum_{h=0}^{2} W_{hj}^{k_2}[\frac{(Z_{k_2}^{(2)}-\nu_j)^2}{2(\tau_j^2)^2} - \frac{1}{2\tau_j^2}]]\}$$

$$= \sum_{k_1 \neq k_2}[\sum_{i=0}^{2}\sum_{h=0}^{2} E[W_{ij}^{k_1}W_{hj}^{k_2}][\frac{(Z_{k_1}^{(2)}-\nu_j)^2}{2(\tau_j^2)^2} - \frac{1}{2\tau_j^2}][\frac{(Z_{k_2}^{(2)}-\nu_j)^2}{2(\tau_j^2)^2} - \frac{1}{2\tau_j^2}]]$$

$$+ \sum_{k_1=k_2=k}[\sum_{i=0}^{2}\sum_{h=0}^{2} E[W_{ij}^{k}W_{hj}^{k}](\frac{(Z_{k}^{(2)}-\nu_j)^2}{2(\tau_j^2)^2} - \frac{1}{2\tau_j^2})^2]$$

As $W_{ij}^k \in \{0,1\}$, when $i = h$ we have $E[W_{ij}^k W_{hj}^k] = E[W_{ij}^k W_{ij}^k] = E[W_{ij}^k]$. What's more, when $i \neq h$ we have $E[W_{ij}^k W_{hj}^k] = 0$ because $W_{ij}^k, W_{hj}^k \in \{0,1\}$ and $W_{ij}^k, W_{hj}^k$ not equal to 1 at the same time. Then we have

$$E[S(\tau_j^2)S(\tau_j^2)]$$

$$= \sum_{k_1 \neq k_2} [\sum_{i=0}^{2} \sum_{h=0}^{2} E[W_{ij}^{k_1}] E[W_{hj}^{k_2}] [\frac{(Z_{k_1}^{(2)} - \nu_j)^2}{2(\tau_j^2)^2} - \frac{1}{2\tau_j^2}][\frac{(Z_{k_2}^{(2)} - \nu_j)^2}{2(\tau_j^2)^2} - \frac{1}{2\tau_j^2}]]$$

$$+ \sum_{k_1 = k_2 = k} [\sum_{i=0}^{2} E[W_{ij}^{k}] (\frac{(Z_{k}^{(2)} - \nu_j)^2}{2(\tau_j^2)^2} - \frac{1}{2\tau_j^2})^2]$$

11)

$$S(\tau_j^2) S(\tau_l^2) = \frac{\partial}{\partial \tau_j^2} \times \frac{\partial}{\partial \tau_l^2} =$$

$$[\sum_{k_1=1}^{N} \sum_{i=0}^{2} W_{ij}^{k_1} [\frac{(Z_{k_1}^{(2)} - \nu_j)^2}{2(\tau_j^2)^2} - \frac{1}{2\tau_j^2}]][\sum_{k_2=1}^{N} \sum_{h=0}^{2} W_{hl}^{k_2} [\frac{(Z_{k_2}^{(2)} - \nu_l)^2}{2(\tau_l^2)^2} - \frac{1}{2\tau_l^2}]]$$

Where $i, h \in \{0, 1, 2\}$ and $j.l \in \{1, 2\}$.

$$E[S(\tau_j^2) S(\tau_l^2)] = E[\frac{\partial}{\partial \tau_j^2} \times \frac{\partial}{\partial \tau_l^2}]$$

$$= E\{[\sum_{k_1=1}^{N} \sum_{i=0}^{2} W_{ij}^{k_1} [\frac{(Z_{k_1}^{(2)} - \nu_j)^2}{2(\tau_j^2)^2} - \frac{1}{2\tau_j^2}]][\sum_{k_2=1}^{N} \sum_{h=0}^{2} W_{hl}^{k_2} [\frac{(Z_{k_2}^{(2)} - \nu_l)^2}{2(\tau_l^2)^2} - \frac{1}{2\tau_l^2}]]\}$$

$$= \sum_{k_1 \neq k_2} [\sum_{i=0}^{2} \sum_{h=0}^{2} E[W_{ij}^{k_1} W_{hl}^{k_2}] [\frac{(Z_{k_1}^{(2)} - \nu_j)^2}{2(\tau_j^2)^2} - \frac{1}{2\tau_j^2}][\frac{(Z_{k_2}^{(2)} - \nu_l)^2}{2(\tau_l^2)^2} - \frac{1}{2\tau_l^2}]]$$

$$+ \sum_{k_1 = k_2 = k} [\sum_{i=0}^{2} \sum_{h=0}^{2} E[W_{ij}^{k} W_{hl}^{k}] [\frac{(Z_{k}^{(2)} - \nu_j)^2}{2(\tau_j^2)^2} - \frac{1}{2\tau_j^2}][\frac{(Z_{k}^{(2)} - \nu_l)^2}{2(\tau_l^2)^2} - \frac{1}{2\tau_l^2}]]$$

As $W_{ij}^k, W_{hl}^k \in \{0, 1\}$, $E[W_{ij}^k W_{hl}^k] = 0$ because $W_{ij}^k, W_{hl}^k$ can not be 1 at the same time. Then we have

$$E[S(\tau_j^2) S(\tau_l^2)]$$

$$= \sum_{k_1 \neq k_2} [\sum_{i=0}^{2} \sum_{h=0}^{2} E[W_{ij}^{k_1}] E[W_{hl}^{k_2}] [\frac{(Z_{k_1}^{(2)} - \nu_j)^2}{2(\tau_j^2)^2} - \frac{1}{2\tau_j^2}][\frac{(Z_{k_2}^{(2)} - \nu_l)^2}{2(\tau_l^2)^2} - \frac{1}{2\tau_l^2}]]$$

12)

$$S(\sigma_i^2) S(\tau_l^2) = \frac{\partial}{\partial \sigma_i^2} \times \frac{\partial}{\partial \tau_l^2} =$$

$$[\sum_{k_1=1}^{N} \sum_{j=0}^{2} W_{ij}^{k_1} [\frac{(Z_{k_1}^{(1)} - \mu_i)^2}{2(\sigma_i^2)^2} - \frac{1}{2\sigma_i^2}]][\sum_{k_2=1}^{N} \sum_{h=0}^{2} W_{hl}^{k_2} [\frac{(Z_{k_2}^{(2)} - \nu_l)^2}{2(\tau_l^2)^2} - \frac{1}{2\tau_l^2}]]$$

Where $i, l \in \{1, 2\}$, $j, h \in \{0, 1, 2\}$.

$$E[S(\sigma_i^2) S(\tau_l^2)] = E[\frac{\partial}{\partial \sigma_i^2} \times \frac{\partial}{\partial \tau_l^2}]$$

$$= E\{[\sum_{k_1=1}^{N}\sum_{j=0}^{2} W_{ij}^{k_1}[\frac{(Z_{k_1}^{(1)}-\mu_i)^2}{2(\sigma_i^2)^2}-\frac{1}{2\sigma_i^2}]][\sum_{k_2=1}^{N}\sum_{h=0}^{2} W_{hl}^{k_2}[\frac{(Z_{k_2}^{(2)}-\nu_l)^2}{2(\tau_l^2)^2}-\frac{1}{2\tau_l^2}]]\}$$

$$= \sum_{k_1\neq k_2}[\sum_{j=0}^{2}\sum_{h=0}^{2} E[W_{ij}^{k_1}W_{hl}^{k_2}][\frac{(Z_{k_1}^{(1)}-\mu_i)^2}{2(\sigma_i^2)^2}-\frac{1}{2\sigma_i^2}][\frac{(Z_{k_2}^{(2)}-\nu_l)^2}{2(\tau_l^2)^2}-\frac{1}{2\tau_l^2}]]$$

$$+ \sum_{k_1=k_2=k}[\sum_{j=0}^{2}\sum_{h=0}^{2} E[W_{ij}^{k}W_{hl}^{k}][\frac{(Z_{k}^{(1)}-\mu_i)^2}{2(\sigma_i^2)^2}-\frac{1}{2\sigma_i^2}][\frac{(Z_{k}^{(2)}-\nu_l)^2}{2(\tau_l^2)^2}-\frac{1}{2\tau_l^2}]]$$

$E[W_{ij}^kW_{hl}^k] = E[W_{il}^k]$ because $W_{ij}^k, W_{hl}^k \in \{0,1\}$ can not be 1 at the same time unless $(i,j) = (h,l) = (i,l)$. Then we have

$$E[S(\sigma_i^2)S(\tau_l^2)]$$

$$= \sum_{k_1\neq k_2}[\sum_{j=0}^{2}\sum_{l=0}^{2} E[W_{ij}^{k_1}]E[W_{hl}^{k_2}][\frac{(Z_{k_1}^{(1)}-\mu_i)^2}{2(\sigma_i^2)^2}-\frac{1}{2\sigma_i^2}][\frac{(Z_{k_2}^{(2)}-\nu_l)^2}{2(\tau_l^2)^2}-\frac{1}{2\tau_l^2}]]$$

$$+ \sum_{k_1=k_2=k}[E[W_{il}^k][\frac{(Z_{k}^{(1)}-\mu_i)^2}{2(\sigma_i^2)^2}-\frac{1}{2\sigma_i^2}][\frac{(Z_{k}^{(2)}-\nu_l)^2}{2(\tau_l^2)^2}-\frac{1}{2\tau_l^2}]]$$

13)

$$S(\mu_i)S(\sigma_i^2) = \frac{\partial}{\partial\mu_i}\times\frac{\partial}{\partial\sigma_i^2} =$$

$$[\sum_{k_1=1}^{N}\sum_{j=0}^{2} W_{ij}^{k_1}\frac{(Z_{k_1}^{(1)}-\mu_i)}{\sigma_i^2}][\sum_{k_2=1}^{N}\sum_{l=0}^{2} W_{il}^{k_2}[\frac{(Z_{k_2}^{(1)}-\mu_i)^2}{2(\sigma_i^2)^2}-\frac{1}{2\sigma_i^2}]]$$

Where $i \in \{1,2\}$, $j,l \in \{0,1,2\}$.

$$E[S(\mu_i)S(\sigma_i^2)] = E[\frac{\partial}{\partial\mu_i}\times\frac{\partial}{\partial\sigma_i^2}]$$

$$= E\{[\sum_{k_1=1}^{N}\sum_{j=0}^{2} W_{ij}^{k_1}\frac{(Z_{k_1}^{(1)}-\mu_i)}{\sigma_i^2}][\sum_{k_2=1}^{N}\sum_{l=0}^{2} W_{il}^{k_2}[\frac{(Z_{k_2}^{(1)}-\mu_i)^2}{2(\sigma_i^2)^2}-\frac{1}{2\sigma_i^2}]]\}$$

$$= \sum_{k_1\neq k_2}[\sum_{j=0}^{2}\sum_{l=0}^{2} E[W_{ij}^{k_1}W_{il}^{k_2}][\frac{(Z_{k_1}^{(1)}-\mu_i)}{\sigma_i^2}][\frac{(Z_{k_2}^{(1)}-\mu_i)^2}{2(\sigma_i^2)^2}-\frac{1}{2\sigma_i^2}]]$$

$$+ \sum_{k_1=k_2=k}[\sum_{j=0}^{2}\sum_{l=0}^{2} E[W_{ij}^{k}W_{il}^{k}][\frac{(Z_{k}^{(1)}-\mu_i)}{\sigma_i^2}][\frac{(Z_{k}^{(1)}-\mu_i)^2}{2(\sigma_i^2)^2}-\frac{1}{2\sigma_i^2}]]$$

$E[W_{ij}^kW_{il}^k] = E[W_{ij}^k]$ because $W_{ij}^k, W_{il}^k \in \{0,1\}$ can not be 1 at the same time unless $j = l$. Then we have

$$E[S(\mu_i)S(\sigma_i^2)]$$

$$= \sum_{k_1 \neq k_2} [\sum_{j=0}^{2} \sum_{l=0}^{2} E[W_{ij}^{k_1}] E[W_{hl}^{k_2}] [\frac{(Z_{k_1}^{(1)} - \mu_i)}{\sigma_i^2}] [\frac{(Z_{k_2}^{(1)} - \mu_i)^2}{2(\sigma_i^2)^2} - \frac{1}{2\sigma_i^2}]]$$

$$+ \sum_{k_1 = k_2 = k} [\sum_{j=0}^{2} E[W_{ij}^{k}] [\frac{(Z_{k}^{(1)} - \mu_i)}{\sigma_i^2}] [\frac{(Z_{k}^{(1)} - \mu_i)^2}{2(\sigma_i^2)^2} - \frac{1}{2\sigma_i^2}]]$$

14)

$$S(\mu_i) S(\sigma_h^2) = \frac{\partial}{\partial \mu_i} \times \frac{\partial}{\partial \sigma_h^2} =$$

$$[\sum_{k_1=1}^{N} \sum_{j=0}^{2} W_{ij}^{k_1} \frac{(Z_{k_1}^{(1)} - \mu_i)}{\sigma_i^2}] [\sum_{k_2=1}^{N} \sum_{l=0}^{2} W_{hl}^{k_2} [\frac{(Z_{k_2}^{(1)} - \mu_h)^2}{2(\sigma_h^2)^2} - \frac{1}{2\sigma_h^2}]]$$

Where $i, h \in \{1, 2\}$, $j, l \in \{0, 1, 2\}$ and $i \neq h$.

$$E[S(\mu_i) S(\sigma_h^2)] = E[\frac{\partial}{\partial \mu_i} \times \frac{\partial}{\partial \sigma_h^2}]$$

$$= E\{[\sum_{k_1=1}^{N} \sum_{j=0}^{2} W_{ij}^{k_1} \frac{(Z_{k_1}^{(1)} - \mu_i)}{\sigma_i^2}] [\sum_{k_2=1}^{N} \sum_{l=0}^{2} W_{hl}^{k_2} [\frac{(Z_{k_2}^{(1)} - \mu_h)^2}{2(\sigma_h^2)^2} - \frac{1}{2\sigma_h^2}]]\}$$

$$= \sum_{k_1 \neq k_2} [\sum_{j=0}^{2} \sum_{l=0}^{2} E[W_{ij}^{k_1} W_{hl}^{k_2}] [\frac{(Z_{k_1}^{(1)} - \mu_i)}{\sigma_i^2}] [\frac{(Z_{k_2}^{(1)} - \mu_h)^2}{2(\sigma_h^2)^2} - \frac{1}{2\sigma_h^2}]]$$

$$+ \sum_{k_1 = k_2 = k} [\sum_{j=0}^{2} \sum_{l=0}^{2} E[W_{ij}^{k} W_{hl}^{k}] [\frac{(Z_{k}^{(1)} - \mu_i)}{\sigma_i^2}] [\frac{(Z_{k}^{(1)} - \mu_h)^2}{2(\sigma_h^2)^2} - \frac{1}{2\sigma_h^2}]]$$

$E[W_{ij}^{k} W_{hl}^{k}] = 0$ because $W_{ij}^{k}, W_{hl}^{k} \in \{0, 1\}$ can not be 1 at the same time. Then we have

$$E[S(\mu_i) S(\sigma_h^2)]$$

$$= \sum_{k_1 \neq k_2} [\sum_{j=0}^{2} \sum_{l=0}^{2} E[W_{ij}^{k_1}] E[W_{hl}^{k_2}] [\frac{(Z_{k_1}^{(1)} - \mu_i)}{\sigma_i^2}] [\frac{(Z_{k_2}^{(1)} - \mu_h)^2}{2(\sigma_h^2)^2} - \frac{1}{2\sigma_h^2}]]$$

15)

$$S(\nu_j) S(\tau_j^2) = \frac{\partial}{\partial \nu_j} \times \frac{\partial}{\partial \tau_j^2} =$$

$$[\sum_{k_1=1}^{N} \sum_{i=0}^{2} W_{ij}^{k_1} \frac{(Z_{k_1}^{(2)} - \nu_j)}{\tau_j^2}] [\sum_{k_2=1}^{N} \sum_{h=0}^{2} W_{hj}^{k_2} [\frac{(Z_{k_2}^{(2)} - \nu_j)^2}{2(\tau_j^2)^2} - \frac{1}{2\tau_j^2}]]$$

Where $i, h \in \{0, 1, 2\}$, $j \in \{1, 2\}$.

$$E[S(\nu_j) S(\tau_j^2)] = E[\frac{\partial}{\partial \nu_j} \times \frac{\partial}{\partial \tau_j^2}]$$

$$= E\{[\sum_{k_1=1}^{N}\sum_{i=0}^{2} W_{ij}^{k_1}\frac{(Z_{k_1}^{(2)}-\nu_j)}{\tau_j^2}][\sum_{k_2=1}^{N}\sum_{h=0}^{2} W_{hj}^{k_2}[\frac{(Z_{k_2}^{(2)}-\nu_j)^2}{2(\tau_j^2)^2}-\frac{1}{2\tau_j^2}]]\}$$

$$= \sum_{k_1\neq k_2}[\sum_{i=0}^{2}\sum_{h=0}^{2} E[W_{ij}^{k_1}W_{hj}^{k_2}][\frac{(Z_{k_1}^{(2)}-\nu_j)}{\tau_j^2}][\frac{(Z_{k_2}^{(2)}-\nu_j)^2}{2(\tau_j^2)^2}-\frac{1}{2\tau_j^2}]]$$

$$+ \sum_{k_1=k_2=k}[\sum_{i=0}^{2}\sum_{h=0}^{2} E[W_{ij}^{k}W_{hj}^{k}][\frac{(Z_{k}^{(2)}-\nu_j)}{\tau_j^2}][\frac{(Z_{k}^{(2)}-\nu_j)^2}{2(\tau_j^2)^2}-\frac{1}{2\tau_j^2}]]$$

$E[W_{ij}^{k}W_{hj}^{k}] = E[W_{ij}^{k}]$ because $W_{ij}^{k}, W_{hl}^{k} \in \{0,1\}$ can not be 1 at the same time unless $(i,j)=(h,l)$. Then we have

$$E[S(\nu_j)S(\tau_j^2)]$$

$$= \sum_{k_1\neq k_2}[\sum_{i=0}^{2}\sum_{h=0}^{2} E[W_{ij}^{k_1}]E[W_{hj}^{k_2}][\frac{(Z_{k_1}^{(2)}-\nu_j)}{\tau_j^2}][\frac{(Z_{k_2}^{(2)}-\nu_j)^2}{2(\tau_j^2)^2}-\frac{1}{2\tau_j^2}]]$$

$$+ \sum_{k_1=k_2=k}[\sum_{i=0}^{2} E[W_{ij}^{k}][\frac{(Z_{k}^{(2)}-\nu_j)}{\tau_j^2}][\frac{(Z_{k}^{(2)}-\nu_j)^2}{2(\tau_j^2)^2}-\frac{1}{2\tau_j^2}]]$$

16)

$$S(\nu_j)S(\tau_l^2) = \frac{\partial}{\partial\nu_j}\times\frac{\partial}{\partial\tau_l^2} =$$

$$[\sum_{k_1=1}^{N}\sum_{i=0}^{2} W_{ij}^{k_1}\frac{(Z_{k_1}^{(2)}-\nu_j)}{\tau_j^2}][\sum_{k_2=1}^{N}\sum_{h=0}^{2} W_{hl}^{k_2}[\frac{(Z_{k_2}^{(2)}-\nu_l)^2}{2(\tau_l^2)^2}-\frac{1}{2\tau_l^2}]]$$

Where $i,h \in \{0,1,2\}$, $j,l \in \{1,2\}$ and $j \neq l$.

$$E[S(\nu_j)S(\tau_l^2)] = E[\frac{\partial}{\partial\nu_j}\times\frac{\partial}{\partial\tau_l^2}]$$

$$= E\{[\sum_{k_1=1}^{N}\sum_{i=0}^{2} W_{ij}^{k_1}\frac{(Z_{k_1}^{(2)}-\nu_j)}{\tau_j^2}][\sum_{k_2=1}^{N}\sum_{h=0}^{2} W_{hl}^{k_2}[\frac{(Z_{k_2}^{(2)}-\nu_l)^2}{2(\tau_l^2)^2}-\frac{1}{2\tau_l^2}]]\}$$

$$= \sum_{k_1\neq k_2}[\sum_{i=0}^{2}\sum_{h=0}^{2} E[W_{ij}^{k_1}W_{hl}^{k_2}][\frac{(Z_{k_1}^{(2)}-\nu_j)}{\tau_j^2}][\frac{(Z_{k_2}^{(2)}-\nu_l)^2}{2(\tau_l^2)^2}-\frac{1}{2\tau_l^2}]]$$

$$+ \sum_{k_1=k_2=k}[\sum_{i=0}^{2}\sum_{h=0}^{2} E[W_{ij}^{k}W_{hl}^{k}][\frac{(Z_{k}^{(2)}-\nu_j)}{\tau_j^2}][\frac{(Z_{k}^{(2)}-\nu_l)^2}{2(\tau_l^2)^2}-\frac{1}{2\tau_l^2}]]$$

$E[W_{ij}^{k}W_{hl}^{k}] = 0$ because $W_{ij}^{k}, W_{hl}^{k} \in \{0,1\}$ can not be 1 at the same time. Then we have

$$E[S(\nu_j)S(\tau_l^2)]$$

$$= \sum_{k_1\neq k_2}[\sum_{i=0}^{2}\sum_{h=0}^{2} E[W_{ij}^{k_1}]E[W_{hl}^{k_2}][\frac{(Z_{k_1}^{(2)}-\nu_j)}{\tau_j^2}][\frac{(Z_{k_2}^{(2)}-\nu_l)^2}{2(\tau_l^2)^2}-\frac{1}{2\tau_l^2}]]$$

17)

$$S(\mu_i)S(\tau_l^2) = \frac{\partial}{\partial \mu_i} \times \frac{\partial}{\partial \tau_l^2} =$$

$$[\sum_{k_1=1}^{N}\sum_{j=0}^{2} W_{ij}^{k_1}\frac{(Z_{k_1}^{(1)}-\mu_i)}{\sigma_i^2}][\sum_{k_2=1}^{N}\sum_{h=0}^{2} W_{hl}^{k_2}[\frac{(Z_{k_2}^{(2)}-\nu_l)^2}{2(\tau_l^2)^2} - \frac{1}{2\tau_l^2}]]$$

Where $i, l \in \{1,2\}$, $j, h \in \{0,1,2\}$.

$$E[S(\mu_i)S(\tau_l^2)] = E[\frac{\partial}{\partial \mu_i} \times \frac{\partial}{\partial \tau_l^2}]$$

$$= E\{[\sum_{k_1=1}^{N}\sum_{j=0}^{2} W_{ij}^{k_1}\frac{(Z_{k_1}^{(1)}-\mu_i)}{\sigma_i^2}][\sum_{k_2=1}^{N}\sum_{h=0}^{2} W_{hl}^{k_2}[\frac{(Z_{k_2}^{(2)}-\nu_l)^2}{2(\tau_l^2)^2} - \frac{1}{2\tau_l^2}]]\}$$

$$= \sum_{k_1\neq k_2}[\sum_{j=0}^{2}\sum_{h=0}^{2} E[W_{ij}^{k_1}W_{hl}^{k_2}][\frac{(Z_{k_1}^{(1)}-\mu_i)}{\sigma_i^2}][\frac{(Z_{k_2}^{(2)}-\nu_l)^2}{2(\tau_l^2)^2} - \frac{1}{2\tau_l^2}]]$$

$$+ \sum_{k_1=k_2=k}[\sum_{j=0}^{2}\sum_{h=0}^{2} E[W_{ij}^{k}W_{hl}^{k}][\frac{(Z_{k}^{(1)}-\mu_i)}{\sigma_i^2}][\frac{(Z_{k}^{(2)}-\nu_l)^2}{2(\tau_l^2)^2} - \frac{1}{2\tau_l^2}]]$$

$E[W_{ij}^k W_{hl}^k] = E[W_{il}^k]$ because $W_{ij}^k, W_{hl}^k \in \{0,1\}$ can not be 1 at the same time unless $(i,j) = (h,l) = (i,l)$. Then we have

$$E[S(\mu_i)S(\tau_l^2)]$$

$$= \sum_{k_1\neq k_2}[\sum_{j=0}^{2}\sum_{h=0}^{2} E[W_{ij}^{k_1}]E[W_{hl}^{k_2}][\frac{(Z_{k_1}^{(1)}-\mu_i)}{\sigma_i^2}][\frac{(Z_{k_2}^{(2)}-\nu_l)^2}{2(\tau_l^2)^2} - \frac{1}{2\tau_l^2}]]$$

$$+ \sum_{k_1=k_2=k}[E[W_{il}^{k}][\frac{(Z_{k}^{(1)}-\mu_i)}{\sigma_i^2}][\frac{(Z_{k}^{(2)}-\nu_l)^2}{2(\tau_l^2)^2} - \frac{1}{2\tau_l^2}]]$$

18)

$$S(\nu_j)S(\sigma_h^2) = \frac{\partial}{\partial \nu_j} \times \frac{\partial}{\partial \sigma_h^2} =$$

$$[\sum_{k_1=1}^{N}\sum_{i=0}^{2} W_{ij}^{k_1}\frac{(Z_{k_1}^{(2)}-\nu_j)}{\tau_j^2}][\sum_{k_2=1}^{N}\sum_{l=0}^{2} W_{hl}^{k_2}[\frac{(Z_{k_2}^{(1)}-\mu_h)^2}{2(\sigma_h^2)^2} - \frac{1}{2\sigma_h^2}]]$$

Where $i, l \in \{0,1,2\}$, $j, h \in \{1,2\}$.

$$E[S(\nu_j)S(\sigma_h^2)] = E[\frac{\partial}{\partial \nu_j} \times \frac{\partial}{\partial \sigma_h^2}]$$

$$= E\{[\sum_{k_1=1}^{N}\sum_{i=0}^{2} W_{ij}^{k_1}\frac{(Z_{k_1}^{(2)}-\nu_j)}{\tau_j^2}][\sum_{k_2=1}^{N}\sum_{l=0}^{2} W_{hl}^{k_2}[\frac{(Z_{k_2}^{(1)}-\mu_h)^2}{2(\sigma_h^2)^2} - \frac{1}{2\sigma_h^2}]]\}$$

$$= \sum_{k_1 \neq k_2} [\sum_{i=0}^{2} \sum_{l=0}^{2} E[W_{ij}^{k_1} W_{hl}^{k_2}][\frac{(Z_{k_1}^{(2)} - \nu_j)}{\tau_j^2}][\frac{(Z_{k_2}^{(1)} - \mu_h)^2}{2(\sigma_h^2)^2} - \frac{1}{2\sigma_h^2}]]$$

$$+ \sum_{k_1 = k_2 = k} [\sum_{i=0}^{2} \sum_{l=0}^{2} E[W_{ij}^{k} W_{hl}^{k}][\frac{(Z_{k}^{(2)} - \nu_j)}{\tau_j^2}][\frac{(Z_{k}^{(1)} - \mu_h)^2}{2(\sigma_h^2)^2} - \frac{1}{2\sigma_h^2}]]$$

$E[W_{ij}^k W_{hl}^k] = E[W_{hj}^k]$ because $W_{ij}^k, W_{hl}^k \in \{0, 1\}$ can not be 1 at the same time unless $(i, j) = (h, l) = (h, j)$. Then we have

$$E[S(\nu_j) S(\sigma_h^2)]$$

$$= \sum_{k_1 \neq k_2} [\sum_{i=0}^{2} \sum_{l=0}^{2} E[W_{ij}^{k_1}] E[W_{hl}^{k_2}][\frac{(Z_{k_1}^{(2)} - \nu_j)}{\tau_j^2}][\frac{(Z_{k_2}^{(1)} - \mu_h)^2}{2(\sigma_h^2)^2} - \frac{1}{2\sigma_h^2}]]$$

$$+ \sum_{k_1 = k_2 = k} [E[W_{hj}^{k}][\frac{(Z_{k}^{(2)} - \nu_j)}{\tau_j^2}][\frac{(Z_{k}^{(1)} - \mu_h)^2}{2(\sigma_h^2)^2} - \frac{1}{2\sigma_h^2}]]$$

19)

$$S(\pi_{ij}) S(\mu_i) = \frac{\partial}{\partial \pi_{ij}} \times \frac{\partial}{\partial \mu_i} =$$

$$[\sum_{k_1=1}^{N} (\frac{W_{ij}^{k_1}}{\pi_{ij}} - \frac{W_{00}^{k_1}}{\pi_{00}})][\sum_{k_2=1}^{N} \sum_{l=0}^{2} W_{il}^{k_2} \frac{Z_{k_2}^{(1)} - \mu_i}{\sigma_i^2}]$$

Where $i \in \{1, 2\}$, $j, l \in \{0, 1, 2\}$ and $i + j \neq 0$.

$$E[S(\pi_{ij}) S(\mu_i)] = E[\frac{\partial}{\partial \pi_{ij}} \times \frac{\partial}{\partial \mu_i}]$$

$$= E\{[\sum_{k_1=1}^{N} (\frac{W_{ij}^{k_1}}{\pi_{ij}} - \frac{W_{00}^{k_1}}{\pi_{00}})][\sum_{k_2=1}^{N} \sum_{l=0}^{2} W_{il}^{k_2} \frac{Z_{k_2}^{(1)} - \mu_i}{\sigma_i^2}]\}$$

$$= \sum_{k_1 \neq k_2} [\sum_{l=0}^{2} (\frac{E[W_{il}^{k_2} W_{ij}^{k_1}]}{\pi_{ij}} - \frac{E[W_{il}^{k_2} W_{00}^{k_1}]}{\pi_{00}}) \frac{Z_{k_2}^{(1)} - \mu_i}{\sigma_i^2}]$$

$$+ \sum_{k_1 = k_2 = k} [\sum_{l=0}^{2} (\frac{E[W_{il}^{k} W_{ij}^{k}]}{\pi_{ij}} - \frac{E[W_{il}^{k} W_{00}^{k}]}{\pi_{00}}) \frac{Z_{k}^{(1)} - \mu_i}{\sigma_i^2}]$$

$E[W_{il}^k W_{ij}^k] = E[W_{ij}^k]$ because $W_{ij}^k, W_{il}^k \in \{0, 1\}$ can not be 1 at the same time unless $(i, j) = (i, l)$. What's more, $E[W_{il}^k W_{00}^k] = 0$ because $W_{il}^k$ and $W_{00}^k$ can not be 1 at the same time. Then we have

$$S(\pi_{ij}) S(\mu_i)$$

$$= \sum_{k_1 \neq k_2} [\sum_{l=0}^{2} (\frac{E[W_{il}^{k_2}] E[W_{ij}^{k_1}]}{\pi_{ij}} - \frac{E[W_{il}^{k_2}] E[W_{00}^{k_1}]}{\pi_{00}}) \frac{Z_{k_2}^{(1)} - \mu_i}{\sigma_i^2}]$$

$$+\sum_{k_1=k_2=k}[\frac{E[W_{ij}^k]}{\pi_{ij}}\frac{Z_k^{(1)}-\mu_i}{\sigma_i^2}]$$

20)

$$S(\pi_{ij})S(\mu_h)=\frac{\partial}{\partial\pi_{ij}}\times\frac{\partial}{\partial\mu_h}=$$

$$[\sum_{k_1=1}^{N}(\frac{W_{ij}^{k_1}}{\pi_{ij}}-\frac{W_{00}^{k_1}}{\pi_{00}})][\sum_{k_2=1}^{N}\sum_{l=0}^{2}W_{hl}^{k_2}\frac{Z_{k_2}^{(1)}-\mu_h}{\sigma_h^2}]$$

Where $h\in\{1,2\}$, $i,j,l\in\{0,1,2\}$, $i\neq h$ and $i+j\neq 0$.

$$E[S(\pi_{ij})S(\mu_h)]=E[\frac{\partial}{\partial\pi_{ij}}\times\frac{\partial}{\partial\mu_h}]$$

$$=E\{[\sum_{k_1=1}^{N}(\frac{W_{ij}^{k_1}}{\pi_{ij}}-\frac{W_{00}^{k_1}}{\pi_{00}})][\sum_{k_2=1}^{N}\sum_{l=0}^{2}W_{hl}^{k_2}\frac{Z_{k_2}^{(1)}-\mu_h}{\sigma_h^2}]\}$$

$$=\sum_{k_1\neq k_2}[\sum_{l=0}^{2}(\frac{E[W_{hl}^{k_2}W_{ij}^{k_1}]}{\pi_{ij}}-\frac{E[W_{hl}^{k_2}W_{00}^{k_1}]}{\pi_{00}})\frac{Z_{k_2}^{(1)}-\mu_h}{\sigma_h^2}]$$

$$+\sum_{k_1=k_2=k}[\sum_{l=0}^{2}(\frac{E[W_{hl}^{k}W_{ij}^{k}]}{\pi_{ij}}-\frac{E[W_{hl}^{k}W_{00}^{k}]}{\pi_{00}})\frac{Z_{k}^{(1)}-\mu_h}{\sigma_h^2}]$$

$E[W_{ij}^kW_{hl}^k]=E[W_{ij}^kW_{00}^k]=0$ because $W_{ij}^k,W_{hl}^k\in\{0,1\}$ and $W_{00}^k$ can not be 1 at the same time. Then we have

$$S(\pi_{ij})S(\mu_h)$$

$$=\sum_{k_1\neq k_2}[\sum_{l=0}^{2}(\frac{E[W_{hl}^{k_2}]E[W_{ij}^{k_1}]}{\pi_{ij}}-\frac{E[W_{hl}^{k_2}]E[W_{00}^{k_1}]}{\pi_{00}})\frac{Z_{k_2}^{(1)}-\mu_h}{\sigma_h^2}]$$

21)

$$S(\pi_{ij})S(\nu_j)=\frac{\partial}{\partial\pi_{ij}}\times\frac{\partial}{\partial\nu_j}=$$

$$[\sum_{k_1=1}^{N}(\frac{W_{ij}^{k_1}}{\pi_{ij}}-\frac{W_{00}^{k_1}}{\pi_{00}})][\sum_{k_2=1}^{N}\sum_{h=0}^{2}W_{hj}^{k_2}\frac{Z_{k_2}^{(2)}-\nu_j}{\tau_j^2}]$$

Where $j\in\{1,2\}$, $i,h\in\{0,1,2\}$ and $i+j\neq 0$.

$$E[S(\pi_{ij})S(\nu_j)]=E[\frac{\partial}{\partial\pi_{ij}}\times\frac{\partial}{\partial\nu_j}]$$

$$=E\{[\sum_{k_1=1}^{N}(\frac{W_{ij}^{k_1}}{\pi_{ij}}-\frac{W_{00}^{k_1}}{\pi_{00}})][\sum_{k_2=1}^{N}\sum_{h=0}^{2}W_{hj}^{k_2}\frac{Z_{k_2}^{(1)}-\nu_j}{\tau_j^2}]\}$$

$$= \sum_{k_1 \neq k_2} [\sum_{h=0}^{2} (\frac{E[W_{hj}^{k_2} W_{ij}^{k_1}]}{\pi_{ij}} - \frac{E[W_{hj}^{k_2} W_{00}^{k_1}]}{\pi_{00}}) \frac{Z_{k_2}^{(2)} - \nu_j}{\tau_j^2}]$$

$$+ \sum_{k_1 = k_2 = k} [\sum_{h=0}^{2} (\frac{E[W_{hj}^{k} W_{ij}^{k}]}{\pi_{ij}} - \frac{E[W_{hj}^{k} W_{00}^{k}]}{\pi_{00}}) \frac{Z_{k}^{(2)} - \nu_j}{\tau_j^2}]$$

$E[W_{hj}^k W_{ij}^k] = E[W_{ij}^k]$ because $W_{ij}^k, W_{hl}^k \in \{0,1\}$ can not be 1 at the same time unless $(i,j) = (h,j)$. What's more, $E[W_{hj}^k W_{00}^k] = 0$ because $W_{hj}^k$ and $W_{00}^k$ can not be 1 at the same time. Then we have

$$S(\pi_{ij})S(\nu_j)$$

$$= \sum_{k_1 \neq k_2} [\sum_{h=0}^{2} (\frac{E[W_{hj}^{k_2}]E[W_{ij}^{k_1}]}{\pi_{ij}} - \frac{E[W_{hj}^{k_2}]E[W_{00}^{k_1}]}{\pi_{00}}) \frac{Z_{k_2}^{(2)} - \nu_j}{\tau_j^2}]$$

$$+ \sum_{k_1 = k_2 = k} [\frac{E[W_{ij}^{k}]}{\pi_{ij}} \frac{Z_{k}^{(2)} - \nu_j}{\tau_j^2}]$$

22)

$$S(\pi_{ij})S(\nu_l) = \frac{\partial}{\partial \pi_{ij}} \times \frac{\partial}{\partial \nu_l} =$$

$$[\sum_{k_1=1}^{N} (\frac{W_{ij}^{k_1}}{\pi_{ij}} - \frac{W_{00}^{k_1}}{\pi_{00}})][\sum_{k_2=1}^{N} \sum_{h=0}^{2} W_{hl}^{k_2} \frac{Z_{k_2}^{(2)} - \nu_l}{\tau_l^2}]$$

Where $l \in \{1,2\}$, $i,j,h \in \{0,1,2\}$, $j \neq l$ and $i + j \neq 0$.

$$E[S(\pi_{ij})S(\nu_l)] = E[\frac{\partial}{\partial \pi_{ij}} \times \frac{\partial}{\partial \nu_l}]$$

$$= E\{[\sum_{k_1=1}^{N} (\frac{W_{ij}^{k_1}}{\pi_{ij}} - \frac{W_{00}^{k_1}}{\pi_{00}})][\sum_{k_2=1}^{N} \sum_{h=0}^{2} W_{hl}^{k_2} \frac{Z_{k_2}^{(1)} - \nu_l}{\tau_l^2}]\}$$

$$= \sum_{k_1 \neq k_2} [\sum_{h=0}^{2} (\frac{E[W_{hl}^{k_2} W_{ij}^{k_1}]}{\pi_{ij}} - \frac{E[W_{hl}^{k_2} W_{00}^{k_1}]}{\pi_{00}}) \frac{Z_{k_2}^{(2)} - \nu_l}{\tau_l^2}]$$

$$+ \sum_{k_1 = k_2 = k} [\sum_{h=0}^{2} (\frac{E[W_{hl}^{k} W_{ij}^{k}]}{\pi_{ij}} - \frac{E[W_{hl}^{k} W_{00}^{k}]}{\pi_{00}}) \frac{Z_{k}^{(2)} - \nu_l}{\tau_l^2}]$$

$E[W_{ij}^k W_{hl}^k] = E[W_{ij}^k W_{00}^k] = 0$ because $W_{ij}^k, W_{hl}^k \in \{0,1\}$ and $W_{00}^k$ can not be 1 at the same time. Then we have

$$S(\pi_{ij})S(\nu_l)$$

$$= \sum_{k_1 \neq k_2} [\sum_{h=0}^{2} (\frac{E[W_{hl}^{k_2}]E[W_{ij}^{k_1}]}{\pi_{ij}} - \frac{E[W_{hl}^{k_2}]E[W_{00}^{k_1}]}{\pi_{00}}) \frac{Z_{k_2}^{(2)} - \nu_l}{\tau_l^2}]$$

23)

$$S(\pi_{ij})S(\sigma_i^2) = \frac{\partial}{\partial \pi_{ij}} \times \frac{\partial}{\partial \sigma_i^2} =$$

$$[\sum_{k_1=1}^{N}(\frac{W_{ij}^{k_1}}{\pi_{ij}} - \frac{W_{00}^{k_1}}{\pi_{00}})][\sum_{k_2=1}^{N}\sum_{l=0}^{2} W_{il}^{k_2}[\frac{(Z_{k_2}^{(1)} - \mu_i)^2}{2(\sigma_i^2)^2} - \frac{1}{2\sigma_i^2}]]$$

Where $i \in \{1,2\}$, $j,l \in \{0,1,2\}$ and $i + j \neq 0$.

$$E[S(\pi_{ij})S(\sigma_i^2)] = E[\frac{\partial}{\partial \pi_{ij}} \times \frac{\partial}{\partial \sigma_i^2}]$$

$$= E\{[\sum_{k_1=1}^{N}(\frac{W_{ij}^{k_1}}{\pi_{ij}} - \frac{W_{00}^{k_1}}{\pi_{00}})][\sum_{k_2=1}^{N}\sum_{l=0}^{2} W_{il}^{k_2}[\frac{(Z_{k_2}^{(1)} - \mu_i)^2}{2(\sigma_i^2)^2} - \frac{1}{2\sigma_i^2}]]\}$$

$$= \sum_{k_1 \neq k_2}[\sum_{l=0}^{2}(\frac{E[W_{il}^{k_2}W_{ij}^{k_1}]}{\pi_{ij}} - \frac{E[W_{il}^{k_2}W_{00}^{k_1}]}{\pi_{00}})[\frac{(Z_{k_2}^{(1)} - \mu_i)^2}{2(\sigma_i^2)^2} - \frac{1}{2\sigma_i^2}]]$$

$$+ \sum_{k_1=k_2=k}[\sum_{l=0}^{2}(\frac{E[W_{il}^{k}W_{ij}^{k}]}{\pi_{ij}} - \frac{E[W_{il}^{k}W_{00}^{k}]}{\pi_{00}})[\frac{(Z_{k}^{(1)} - \mu_i)^2}{2(\sigma_i^2)^2} - \frac{1}{2\sigma_i^2}]]$$

$E[W_{ij}^k W_{il}^k] = E[W_{ij}^k]$ because $W_{ij}^k, W_{il}^k \in \{0,1\}$ can not be 1 at the same time unless $(i,j) = (i,l)$. What's more, $E[W_{il}^k W_{00}^k] = 0$ because $W_{il}^k$ and $W_{00}^k$ can not be 1 at the same time. Then we have

$$S(\pi_{ij})S(\mu_l)$$

$$= \sum_{k_1 \neq k_2}[\sum_{l=0}^{2}(\frac{E[W_{il}^{k_2}]E[W_{ij}^{k_1}]}{\pi_{ij}} - \frac{E[W_{il}^{k_2}]E[W_{00}^{k_1}]}{\pi_{00}})[\frac{(Z_{k_2}^{(1)} - \mu_i)^2}{2(\sigma_i^2)^2} - \frac{1}{2\sigma_i^2}]]$$

$$+ \sum_{k_1=k_2=k}[\frac{E[W_{ij}^{k}]}{\pi_{ij}}[\frac{(Z_{k}^{(1)} - \mu_i)^2}{2(\sigma_i^2)^2} - \frac{1}{2\sigma_i^2}]]$$

24)

$$S(\pi_{ij})S(\sigma_h^2) = \frac{\partial}{\partial \pi_{ij}} \times \frac{\partial}{\partial \sigma_h^2} =$$

$$[\sum_{k_1=1}^{N}(\frac{W_{ij}^{k_1}}{\pi_{ij}} - \frac{W_{00}^{k_1}}{\pi_{00}})][\sum_{k_2=1}^{N}\sum_{l=0}^{2} W_{hl}^{k_2}[\frac{(Z_{k_2}^{(1)} - \mu_h)^2}{2(\sigma_h^2)^2} - \frac{1}{2\sigma_h^2}]]$$

Where $h \in \{1,2\}$, $i,j,l \in \{0,1,2\}$, $i \neq h$ and $i + j \neq 0$.

$$E[S(\pi_{ij})S(\sigma_h^2)] = E[\frac{\partial}{\partial \pi_{ij}} \times \frac{\partial}{\partial \sigma_h^2}]$$

$$= E\{[\sum_{k_1=1}^{N}(\frac{W_{ij}^{k_1}}{\pi_{ij}} - \frac{W_{00}^{k_1}}{\pi_{00}})][\sum_{k_2=1}^{N}\sum_{l=0}^{2} W_{hl}^{k_2}[\frac{(Z_{k_2}^{(1)} - \mu_h)^2}{2(\sigma_h^2)^2} - \frac{1}{2\sigma_h^2}]]\}$$

$$= \sum_{k_1 \neq k_2}[\sum_{l=0}^{2}(\frac{E[W_{hl}^{k_2}W_{ij}^{k_1}]}{\pi_{ij}} - \frac{E[W_{hl}^{k_2}W_{00}^{k_1}]}{\pi_{00}})[\frac{(Z_{k_2}^{(1)} - \mu_h)^2}{2(\sigma_h^2)^2} - \frac{1}{2\sigma_h^2}]]$$

$$+ \sum_{k_1=k_2=k}[\sum_{l=0}^{2}(\frac{E[W_{hl}^{k}W_{ij}^{k}]}{\pi_{ij}} - \frac{E[W_{hl}^{k}W_{00}^{k}]}{\pi_{00}})[\frac{(Z_{k}^{(1)} - \mu_h)^2}{2(\sigma_h^2)^2} - \frac{1}{2\sigma_h^2}]]$$

$E[W_{ij}^k W_{hl}^k] = E[W_{ij}^k W_{00}^k] = 0$ because $W_{ij}^k, W_{hl}^k \in \{0, 1\}$ and $W_{00}^k$ can not be 1 at the same time. Then we have

$$S(\pi_{ij})S(\mu_h)$$

$$= \sum_{k_1 \neq k_2}[\sum_{l=0}^{2}(\frac{E[W_{hl}^{k_2}]E[W_{ij}^{k_1}]}{\pi_{ij}} - \frac{E[W_{hl}^{k_2}]E[W_{00}^{k_1}]}{\pi_{00}})[\frac{(Z_{k_2}^{(1)} - \mu_h)^2}{2(\sigma_h^2)^2} - \frac{1}{2\sigma_h^2}]]$$

25)

$$S(\pi_{ij})S(\tau_j^2) = \frac{\partial}{\partial \pi_{ij}} \times \frac{\partial}{\partial \tau_j^2} =$$

$$[\sum_{k_1=1}^{N}(\frac{W_{ij}^{k_1}}{\pi_{ij}} - \frac{W_{00}^{k_1}}{\pi_{00}})][\sum_{k_2=1}^{N}\sum_{h=0}^{2} W_{hj}^{k_2}[\frac{(Z_{k_2}^{(2)} - \nu_j)^2}{2(\tau_j^2)^2} - \frac{1}{2\tau_j^2}]]$$

Where $j \in \{1, 2\}$, $i, h \in \{0, 1, 2\}$ and $i + j \neq 0$.

$$E[S(\pi_{ij})S(\tau_j^2)] = E[\frac{\partial}{\partial \pi_{ij}} \times \frac{\partial}{\partial \tau_j^2}]$$

$$= E\{[\sum_{k_1=1}^{N}(\frac{W_{ij}^{k_1}}{\pi_{ij}} - \frac{W_{00}^{k_1}}{\pi_{00}})][\sum_{k_2=1}^{N}\sum_{h=0}^{2} W_{hj}^{k_2}[\frac{(Z_{k_2}^{(2)} - \nu_j)^2}{2(\tau_j^2)^2} - \frac{1}{2\tau_j^2}]]\}$$

$$= \sum_{k_1 \neq k_2}[\sum_{h=0}^{2}(\frac{E[W_{hj}^{k_2}W_{ij}^{k_1}]}{\pi_{ij}} - \frac{E[W_{hj}^{k_2}W_{00}^{k_1}]}{\pi_{00}})[\frac{(Z_{k_2}^{(2)} - \nu_j)^2}{2(\tau_j^2)^2} - \frac{1}{2\tau_j^2}]]$$

$$+ \sum_{k_1=k_2=k}[\sum_{h=0}^{2}(\frac{E[W_{hj}^{k}W_{ij}^{k}]}{\pi_{ij}} - \frac{E[W_{hj}^{k}W_{00}^{k}]}{\pi_{00}})[\frac{(Z_{k}^{(2)} - \nu_j)^2}{2(\tau_j^2)^2} - \frac{1}{2\tau_j^2}]]$$

$E[W_{hj}^k W_{ij}^k] = E[W_{ij}^k]$ because $W_{ij}^k, W_{hl}^k \in \{0, 1\}$ can not be 1 at the same time unless $(i, j) = (h, j)$. What's more, $E[W_{hj}^k W_{00}^k] = 0$ because $W_{hj}^k$ and $W_{00}^k$ can not be 1 at the same time. Then we have

$$S(\pi_{ij})S(\tau_j^2)$$

$$= \sum_{k_1 \neq k_2} [\sum_{h=0}^{2} (\frac{E[W_{hj}^{k_2}]E[W_{ij}^{k_1}]}{\pi_{ij}} - \frac{E[W_{hj}^{k_2}]E[W_{00}^{k_1}]}{\pi_{00}})[\frac{(Z_{k_2}^{(2)} - \nu_j)^2}{2(\tau_j^2)^2} - \frac{1}{2\tau_j^2}]]$$

$$+ \sum_{k_1 = k_2 = k} [\frac{E[W_{ij}^{k}]}{\pi_{ij}} [\frac{(Z_{k}^{(2)} - \nu_j)^2}{2(\tau_j^2)^2} - \frac{1}{2\tau_j^2}]]$$

26)

$$S(\pi_{ij})S(\tau_l^2) = \frac{\partial}{\partial \pi_{ij}} \times \frac{\partial}{\partial \tau_l^2} =$$

$$[\sum_{k_1=1}^{N} (\frac{W_{ij}^{k_1}}{\pi_{ij}} - \frac{W_{00}^{k_1}}{\pi_{00}})][\sum_{k_2=1}^{N} \sum_{h=0}^{2} W_{hl}^{k_2} [\frac{(Z_{k_2}^{(2)} - \nu_l)^2}{2(\tau_l^2)^2} - \frac{1}{2\tau_l^2}]]$$

Where $l \in \{1, 2\}$, $i, j, h \in \{0, 1, 2\}$, $j \neq l$ and $i + j \neq 0$.

$$E[S(\pi_{ij})S(\tau_l^2)] = E[\frac{\partial}{\partial \pi_{ij}} \times \frac{\partial}{\partial \tau_l^2}]$$

$$= E\{[\sum_{k_1=1}^{N} (\frac{W_{ij}^{k_1}}{\pi_{ij}} - \frac{W_{00}^{k_1}}{\pi_{00}})][\sum_{k_2=1}^{N} \sum_{h=0}^{2} W_{hl}^{k_2} [\frac{(Z_{k_2}^{(2)} - \nu_l)^2}{2(\tau_l^2)^2} - \frac{1}{2\tau_l^2}]]\}$$

$$= \sum_{k_1 \neq k_2} [\sum_{h=0}^{2} (\frac{E[W_{hl}^{k_2} W_{ij}^{k_1}]}{\pi_{ij}} - \frac{E[W_{hl}^{k_2} W_{00}^{k_1}]}{\pi_{00}})[\frac{(Z_{k_2}^{(2)} - \nu_l)^2}{2(\tau_l^2)^2} - \frac{1}{2\tau_l^2}]]$$

$$+ \sum_{k_1 = k_2 = k} [\sum_{h=0}^{2} (\frac{E[W_{hl}^{k} W_{ij}^{k}]}{\pi_{ij}} - \frac{E[W_{hl}^{k} W_{00}^{k}]}{\pi_{00}})[\frac{(Z_{k}^{(2)} - \nu_l)^2}{2(\tau_l^2)^2} - \frac{1}{2\tau_l^2}]]$$

$E[W_{ij}^k W_{il}^k] = E[W_{ij}^k]$ because $W_{ij}^k, W_{il}^k \in \{0, 1\}$ can not be 1 at the same time unless $(i, j) = (i, l)$. What's more, $E[W_{il}^k W_{00}^k] = 0$ because $W_{il}^k$ and $W_{00}^k$ can not be 1 at the same time. Then we have

$$S(\pi_{ij})S(\tau_l^2)$$

$$= \sum_{k_1 \neq k_2} [\sum_{h=0}^{2} (\frac{E[W_{hl}^{k_2}]E[W_{ij}^{k_1}]}{\pi_{ij}} - \frac{E[W_{hl}^{k_2}]E[W_{00}^{k_1}]}{\pi_{00}})[\frac{(Z_{k_2}^{(2)} - \nu_l)^2}{2(\tau_l^2)^2} - \frac{1}{2\tau_l^2}]]$$

**Part** $\boldsymbol{III\ S^*(X, \theta) S^{*T}(X, \theta)}$

1)

$$S(\pi_{ij}) = \frac{\partial}{\partial \pi_{ij}} = \sum_{k=1}^{N} \{\frac{W_{ij}^k}{\pi_{ij}} - \frac{W_{00}^k}{\pi_{00}}\}$$

$$S^*(\pi_{ij}) = E_\theta\{S(\pi_{ij})\} = \sum_{k=1}^{N} \{\frac{E[W_{ij}^k]}{\pi_{ij}} - \frac{E[W_{00}^k]}{\pi_{00}}\}$$

Where $i \in \{0, 1, 2\}, j \in \{0, 1, 2\}$ and $i + j \neq 0$.

2)

$$S(\mu_i) = \frac{\partial}{\partial \mu_i} = \sum_{k=1}^{N}\sum_{j=0}^{2}\{W_{ij}^k[\frac{Z_k^{(1)} - \mu_i}{\sigma_i^2}]\}$$

$$S^*(\mu_i) = E_\theta\{S(\mu_i)\} = \sum_{k=1}^{N}\sum_{j=0}^{2}\{E[W_{ij}^k][\frac{Z_k^{(1)} - \mu_i}{\sigma_i^2}]\}$$

Where $i \in \{1, 2\}, j \in \{0, 1, 2\}$.

3)

$$S(\nu_j) = \frac{\partial}{\partial \nu_j} = \sum_{k=1}^{N}\sum_{i=0}^{2}\{W_{ij}^k[\frac{Z_k^{(2)} - \nu_j}{\tau_j^2}]\}$$

$$S^*(\nu_j) = E_\theta\{S(\nu_j)\} = \sum_{k=1}^{N}\sum_{i=0}^{2}\{E[W_{ij}^k][\frac{Z_k^{(2)} - \nu_j}{\tau_j^2}]\}$$

Where $i \in \{0, 1, 2\}, j \in \{1, 2\}$.

4)

$$S(\sigma_i^2) = \frac{\partial}{\partial \sigma_i^2} = \sum_{k=1}^{N}\sum_{j=0}^{2}\{W_{ij}^k[\frac{(Z_k^{(1)} - \mu_i)^2}{2(\sigma_i^2)^2} - \frac{1}{2\sigma_i^2}]\}$$

$$S^*(\sigma_i^2) = E_\theta\{S(\sigma_i^2)\} = \sum_{k=1}^{N}\sum_{j=0}^{2}\{E[W_{ij}^k][\frac{(Z_k^{(1)} - \mu_i)^2}{2(\sigma_i^2)^2} - \frac{1}{2\sigma_i^2}]\}$$

Where $i \in \{1, 2\}, j \in \{0, 1, 2\}$.

5)

$$S(\tau_j^2) = \frac{\partial}{\partial \tau_j^2} = \sum_{k=1}^{N}\sum_{i=0}^{2}\{W_{ij}^k[\frac{(Z_k^{(2)} - \nu_j)^2}{2(\tau_j^2)^2} - \frac{1}{2\tau_j^2}]\}$$

$$S^*(\tau_j^2) = E_\theta\{S(\tau_j^2)\} = \sum_{k=1}^{N}\sum_{i=0}^{2}\{E[W_{ij}^k][\frac{(Z_k^{(2)} - \nu_j)^2}{2(\tau_j^2)^2} - \frac{1}{2\tau_j^2}]\}$$

Where $i \in \{0, 1, 2\}, j \in \{1, 2\}$.

When we get the Fisher information matrix $I_Y$ of the mixture model, we can acquired the covariance matrix $V = I_Y^{-1}$. Then we have

$$M^2RI = \frac{\pi_{11} + \pi_{22}}{1 - \pi_{00}}$$

Let

$$\pi_{00} = 1 - \sum_{i+j \neq 0} \pi_{ij}$$

we have

$$M^2RI = \frac{\pi_{11} + \pi_{22}}{\sum_{i+j\neq 0} \pi_{ij}}$$

Then we can calculate the Jacobian matrix of $g(\pi)$ :

$$J_{g(\pi)} = (\frac{\partial g(\pi)}{\partial \pi_{ij}})$$

where $i + j \neq 0$.

$$\frac{\partial g(\pi)}{\partial \pi_{ij}} = \begin{cases} -\frac{\pi_{11}+\pi_{22}}{(\sum_{i+j\neq 0} \pi_{ij})^2} & i \neq j \\ \frac{\sum_{i\neq j} \pi_{ij}}{(\sum_{i+j\neq 0} \pi_{ij})^2} & i = j, i + j \neq 0 \end{cases}$$

By the delta method, we can acquired the variance of $M^2RI$ :

$$\delta^2 = J_{g(\pi)} V J_{g(\pi)}^T$$

### 4. Theoretical confidence interval for $M^2RI$

The 95 confidence interval for $M^2RI$ is

$$(\mu - Z_{0.975}\delta, \mu + Z_{0.975}\delta)$$

Where $\mu$ is the mean value of $M^2RI$ and $Z_{0.975}$ is the upper two-sided normal distribution percentile at level 0.05.
However, when the reproducibility of the association study was close to 1, the upper bound of the theoretical confidence interval might exceed 1. In order to avoid the occurrence of this unreasonable situation, when that happens, we reset the confidence interval

$$(\mu - Z_{0.95}\delta, 1)$$
